# Gut epithelium modifies enteric behaviors during nutritional adversity via distinct peptidergic signaling axes

**DOI:** 10.1101/2025.01.06.631508

**Authors:** Surojit Sural, Zion Walker, Oliver Hobert

## Abstract

Interorgan signaling events are emerging as key regulators of behavioral plasticity. The foregut and hindgut circuits of the *C. elegans* enteric nervous system (ENS) control feeding and defecation behavior, respectively. Here we show that epithelial cells in the midgut integrate feeding state information to control these behavioral outputs by releasing distinct neuropeptidergic signals. In favorable conditions, insulin and non-insulin peptides released from midgut epithelia activate foregut and hindgut enteric neurons, respectively, to sustain normal feeding and defecation behavior. During food scarcity, altered insulin signaling from sensory neurons activates the transcription factor DAF-16/FoxO in midgut epithelia, which blocks both peptidergic signaling axes to the ENS by transcriptionally shutting down the intestinal neuropeptide secretion machinery. Our findings demonstrate that midgut epithelial cells act as integrators to relay internal state information to distinct parts of the enteric nervous system to control animal behavior.

## INTRODUCTION

The enteric nervous system (ENS), also referred to as the ‘second brain’, is an autonomously acting neuronal network present in the digestive tract of all bilaterians, whose primary function is to regulate motility in the alimentary tract ^1–4^. The mammalian ENS consists of millions of neurons that fall into at least 20 neuron types that regulate gut peristalsis required for digestion, absorption and egestion of food contents ^5–7^. Though enteric circuits are able to generate peristaltic behavior independent of CNS input ^4^, factors such as nutritional, reproductive or psychological status of the animal have been shown to influence motility in the gut ^8–10^. Understanding how internal and external factors are sensed and integrated to coordinate the output from enteric circuits will provide novel insights on gastrointestinal physiology and disease.

The ENS in *Caenorhabditis elegans* has been extensively studied in terms of its anatomy, development, molecular topology and behavioral output ^11–19^, which makes it an attractive model for investigating how ENS function is regulated in metazoans. The *C. elegans* ENS consists of 22 neurons, 20 that are present in the foregut (the pharynx), from here on termed as pharyngeal enteric neurons (PENs), and two that innervate the hindgut, from here on termed as hindgut enteric neurons (HENs) (Figure 1A). The 20 PENs are further classified into 14 anatomically and molecularly distinct classes forming a densely interconnected network that controls rhythmic pharyngeal pumping ^13–15^. The two HENs, termed AVL and DVB, are GABAergic neurons that are activated by the neuropeptide NLP-40 released from the intestine, which generates all-or-none action potentials in these neurons to control the rate of defecation ^11,17,19,20^. Rhythmic activation of the HENs, which form an electrically coupled circuit, results in GABA-dependent contraction of enteric muscles leading to expulsion of contents of the gut, a behavior termed as the defecation motor program ^11,21,22^.

**Figure 1.**
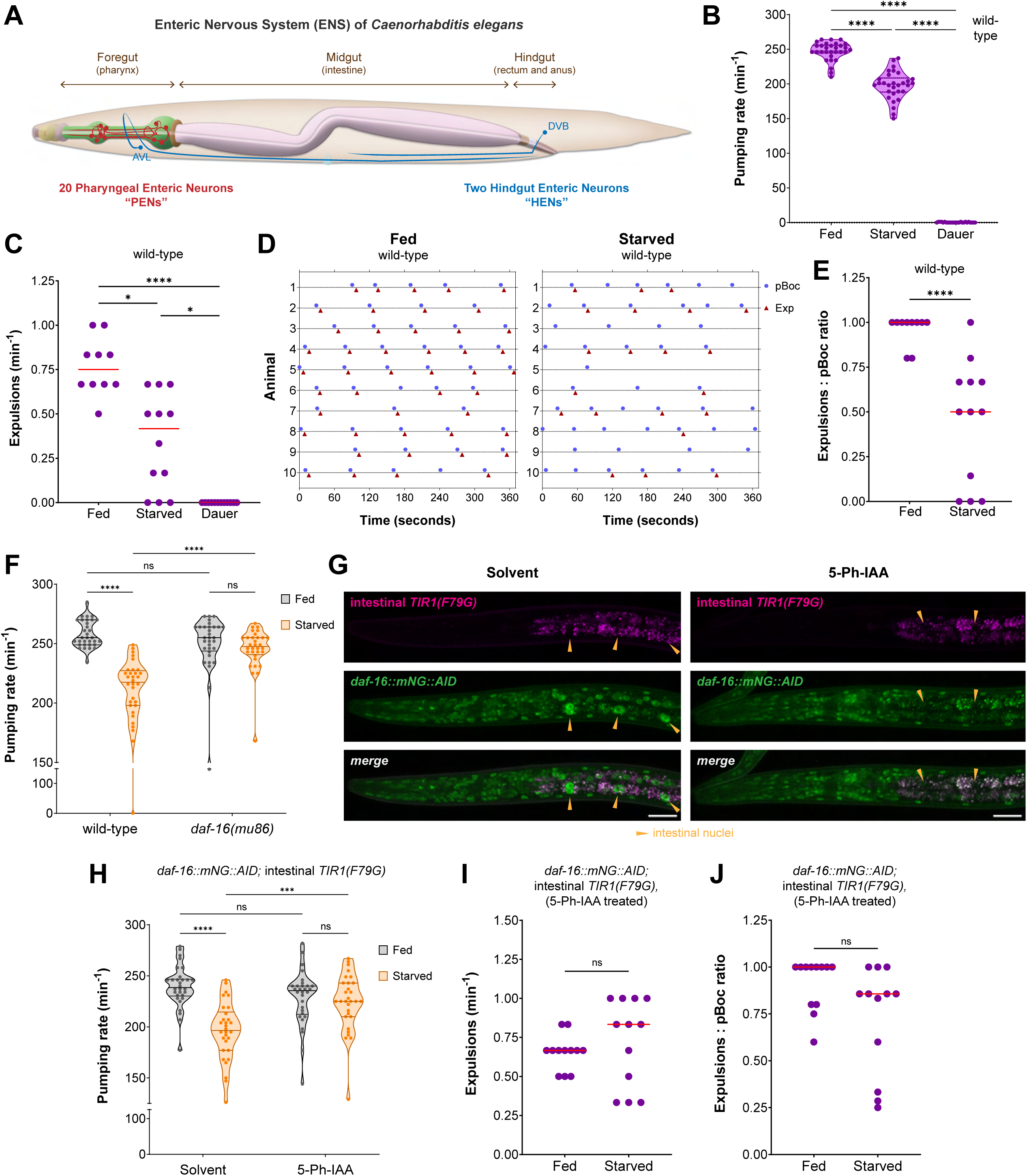
Starvation inhibits enteric nervous system function via cell non-autonomous activity of DAF-16/FoxO in the intestine. (A) Schematic showing the anatomical locations of PENs and HENs in the alimentary tract of *C. elegans*; adapted from WormBook ^98^. (B) Pharyngeal pumping rate on food in fed vs starved (4 hr) wild-type adults and in starvation-induced dauer stage animals. (C) Frequency of expulsion on food in fed vs starved (4 hr) wild-type adults and in starvation-induced dauer stage animals. (D) Representative traces of pBoc and expulsion (Exp) events on food in fed vs starved (4 hr) wild-type adults (10 animals per condition). (E) Expulsion: pBoc ratio on food in fed vs starved (4 hr) wild-type adults. (F) Pharyngeal pumping rate on food in fed vs starved (4 hr) wild-type and *daf-16(mu86)* adults. (G) Intestine-specific DAF-16 depletion in *daf-16(ot853); otSi2[ges-1p::TIR1(F79G)]; daf-2(e1370)* dauer stage animals. Animals were treated with either solvent (ethanol) or 100 μM 5-Ph-IAA. DAF-16 was not detected in the midgut epithelial cells after 5-Ph-IAA treatment in 15/15 animals. *daf-2(e1370)* dauer stage animals were used to visualize nuclear localization of DAF-16 in all tissue types. Scale bars, 20 μm. (H) Pharyngeal pumping rate on food after intestine-specific DAF-16 depletion in fed vs starved (4 hr) *daf-16(ot853); otSi2[ges-1p::TIR1(F79G)]* adults. Animals were treated with either solvent (ethanol) or 100 μM 5-Ph-IAA. (I) Frequency of expulsion on food after intestine-specific DAF-16 depletion in fed vs starved (4 hr) *daf-16(ot853); otSi2[ges-1p::TIR1(F79G)]* adults treated with 100 μM 5-Ph-IAA. (J) Expulsion: pBoc ratio on food after intestine-specific DAF-16 depletion in fed vs starved (4 hr) *daf-16(ot853); otSi2[ges-1p::TIR1(F79G)]* adults treated with 100 μM 5-Ph-IAA.. Horizontal line in the middle of data points represents median value of biological replicates in (B, C, E, F, H-J). Additional horizontal lines represent 25^th^ and 75^th^ percentiles in (B, F, H). *, ***, **** and ns represent *P* < 0.05, *P* < 0.001, *P* < 0.0001 and not significant, respectively, in Dunn’s multiple comparison test after Kruskal-Wallis test in (B, C), in Mann-Whitney test in (E, I, J) and Sidak’s multiple comparisons test after two-way ANOVA in (F, H). See also Figure S1.

After completion of postembryonic development, *C. elegans* continuously feeds and defecates, and the rates of both of these behaviors remain highly consistent both within and across individuals ^22–25^. When an animal is transiently moved away from its food source, there is an immediate reduction in the frequency of foregut and hindgut contractions, which become highly irregular because the animal is no longer actively consuming food ^22–24^. ENS output in the presence of abundant food can also be altered if an animal has experienced starvation in the recent past ^24,26,27^, and this effect is more pronounced in the dauer diapause stage, which the animal enters after experiencing acute nutrient deprivation during early larval development ^28–30^. It remains little understood how different neuronal types in the ENS sense the internal state of an animal to coordinate gut contractility at the systemic level during periods of altered nutritional requirement ^9,31,32^.

Here we show how distinct neuropeptidergic signals are transmitted from midgut epithelial cells to communicate the internal state of the animal to the foregut and hindgut enteric circuits and show that this interorgan signaling is required to maintain normal ENS output in favorable conditions. Previous work has shown that the neuropeptide NLP-40 released from gut epithelia activates the HENs via GPCR signaling to generate normal defecation behavior ^19,20^.

We find that two neuropeptides of the insulin family, INS-7 and INS-35, are also continuously secreted from midgut epithelial cells, and these peptides are sensed by the insulin receptor on neuromodulatory NSM neurons in the foregut to induce rapid foregut contractions in nutrient replete conditions. When an animal undergoes acute starvation, absence of insulin signaling from sensory neurons activates the stress-responsive transcription factor DAF-16/FoxO in the midgut epithelial cells ^33,34^. Using transcriptomic profiling, cell type-specific genetic manipulations and CRISPR/Cas9-generated endogenous reporters, we show that DAF-16/FoxO activation in the midgut epithelia reduces the secretion of neuropeptides from the midgut by transcriptionally silencing not only insulin genes, but also genes that constitute the neuropeptide secretion machinery of the gut epithelial cells. Due to absence of a neuropeptide relay signaling via the intestine to the two enteric circuits, ENS output is greatly reduced both in terms of foregut and hindgut contractions even after the animal returns to a favorable environment after experiencing starvation. Our findings highlight the role of the intestinal epithelia as an integrator of the metabolic status of the animal and unveils the mechanisms of how the non-cell autonomous activity of a stress-responsive transcription factor modifies enteric behaviors by disrupting gut-to-ENS signaling during periods of nutritional adversity.

## RESULTS

### Starvation inhibits output from the ENS via non-cell autonomous activity of DAF-16/FoxO in gut epithelial cells

To study how the nutritional state of an organism alters output from the ENS, we measured how the two enteric behaviors in *C. elegans*, feeding and defecation, are affected by starvation. The rate of feeding, controlled by PENs, is measured as the frequency of foregut contractions, while the rate of defecation, controlled by HENs, is measured as the frequency of expulsion events from the hindgut. The defecation motor program consists of three steps: contraction of posterior body muscles (pBoc), contraction of anterior body muscles (aBoc), and explusion (Exp)^22^. The pBoc step is initiated by a pacemaker in the intestinal epithelial cells ^35^, which acts upstream of HENs, while the Exp step requires activation of the two HENs that constrict hindgut enteric muscles to release the contents of the gut through the anal opening ^19,20^. We measured how feeding and defecation behaviors were affected by starvation after the animal was transferred back to food and allowed an acclimation period of 5 minutes. We used two different starvation paradigms: (1) a four hour period of total food deprivation during adulthood that is sufficient to induce risk-taking behavior during food search ^36^, and (2) food deprivation combined with animal crowding during early development that induces remodeling into the dauer stage of diapause ^28,29^.

We found that starvation during adulthood reduces the rate of feeding even after the animal is returned to food (Figures 1B and S1A). In contrast, animals in the dauer stage display no foregut (pharynx) contractions on food, except for a few sporadic pharyngeal pumps, as described previously (Figure 1B) ^28,30^. We observe a similar effect of starvation on the rate of defecation. The frequency of pBoc events was unaffected after starvation (Figure S1B), but the rate of expulsions was significantly reduced (Figures 1C and 1D). Half of the pBoc events in starved adults did not result in any expulsion of gut contents (Figures 1D and 1E), which indicates altered communication between the intestinal pacemaker and the HENs. Even though these animals underwent a period of food deprivation prior to the measurement of defecation behavior, their gut is filled with food contents in the five minutes of acclimation prior to behavioral recordings, which can be observed during the expulsion events. In contrast, a dauer stage animal does not initiate any pBoc or Exp events when returned to food (Figures 1C and S1B), which is likely attributable to the fact that these animals are not actively feeding because their buccal cavity is sealed ^37^.

Starvation during adulthood or in dauer-inducing conditions during postembryonic development both involve activation of the stress-responsive protein DAF-16 ^29,38,39^, which is the sole *C. elegans* homolog of mammalian FoxO transcription factors. In nutrient replete conditions, agonist insulin peptides signal via DAF-2, the sole member of the insulin/IGF1 receptor family in *C. elegans*, to trigger a phosphorylation cascade that retains DAF-16/FoxO in the cytoplasm in an inactive state (Figure S1C) ^34,40–42^. When the environment becomes unfavorable, absence of agonist insulins and presence of antagonistic insulins inhibit signaling via the DAF-2/InsR receptor and this results in nuclear translocation of unphosphorylated DAF-16/FoxO, where it activates the transcription of stress-responsive genes (Figure S1C) ^34,41,42^. We tested whether starvation affects enteric functions in animals that lack the DAF-16/FoxO transcription factor. We found that *daf-16* null adults do not reduce their rate of feeding when placed on food after the same period of starvation that reduces foregut contractions in wild-type animals (Figure 1F). Similarly, in contrast to wild-type adults (Figures 1C-1E), a 4 hour starvation period did not significantly alter the rate of defecation in adults that lack *daf-16* (Figures S1D-S1F).

A previous study has shown that in the dauer diapause stage, removal of DAF-16/FoxO from the midgut epithelial cells can initiate foregut contractions in conditions that completely silence ENS output ^39^, which suggests a non-cell autonomous role of this transcription factor in regulating ENS function. By visualizing an endogenously GFP-tagged DAF-16/FoxO protein, we found that this transcription factor localizes to the nuclei of midgut epithelial cells in starvation conditions that reduce the rates of feeding and defecation during adulthood (Figure S1G). To test whether removing DAF-16/FoxO only from the midgut epithelial cells can suppress the effects of starvation on enteric behaviors in adults, we generated a single-copy *TIR1(F79G)* transgene driven by the *ges-1* promoter that is only expressed in intestinal epithelial cells (Figure 1G). In a genetic background where *daf-16* is endogenously tagged with AID*, the *TIR1(F79G)* transgene depletes DAF-16/FoxO specifically from the midgut in the presence of 5-Ph-IAA, the synthetic auxin ligand for TIR1(F79G) (Figure 1G) ^43–45^). We found that if DAF-16/FoxO is selectively depleted from gut epithelia, the effect of starvation on feeding rate can be completely suppressed (Figure 1H). Similarly, starvation does not affect defecation behavior in animals that have DAF-16/FoxO depleted only from their midgut epithelial cells (Figures 1I and 1J, and S1H). Unlike the effect of starvation on defecation behavior in wild-type animals (Figures 1D and 1E), DAF-16/FoxO removal specifically from midgut epithelia resulted in the expulsion of gut contents immediately after most pBoc events even in starved animals (Figures 1J and S1H). This establishes a non-cell autonomous role of DAF-16/FoxO to modify output from PENs and HENs to regulate the rates of feeding and defecation, respectively, after a period of nutrient deprivation during adulthood.

### Evidence for two signaling axes from midgut epithelial cells to the ENS

Our previous work has shown that DAF-16/FoxO also functions cell autonomously in the PENs to silence foregut contractions during acute starvation ^39^. This suggests that when food is abundant, insulin peptides released from midgut epithelial cells might act on the PENs via interorgan signaling to prevent DAF-16/FoxO activation and generate normal feeding behavior; while during food deprivation, DAF-16/FoxO activity in the midgut likely alters the release of such gut-derived insulin peptides to activate DAF-16/FoxO in PENs via ‘FoxO-to-FoxO’ signaling ^46^. To test whether enteric neurons actively need insulin signaling to maintain normal ENS output in well-fed conditions, we inhibited DAF-2/InsR specifically in all enteric neurons by expressing a dominant negative form of the DAF-2 receptor, using a driver that is expressed in both PENs and HENs (Figure S2A)^39^. This dominant negative form of the protein, termed DAF-2(DN), has the intracellular kinase domain replaced by a fluorescent protein (Figure S2A), which allows DAF-2(DN) to bind insulin peptides on the extracellular side but does not allow the signal to be transmitted to its intracellular downstream effectors. Insulin/IGF1 receptors need to be phosphorylated by their dimerization partner to achieve activation ^47^, but the lack of a kinase domain in DAF-2(DN) results in inactivation of the wild-type DAF-2/InsR molecule it dimerizes with and thus producing a dominant negative effect.

We found that expression of DAF-2(DN) in all neurons of the ENS (HENs and PENs) is sufficient to reduce the animal’s rate of feeding even in nutrient-rich conditions, but it has no effect on defecation behavior (Figures 2A and S2B-S2E). In terms of change in the rate of feeding, the effect of inhibiting DAF-2/InsR signaling in PENs of well-fed animals is identical to the effect of starvation on wild-type animals (Figure 2A). When animals expressing DAF-2(DN) in PENs were subjected to starvation, it does not further reduce their rate of feeding (Figure 2A), which indicates that starvation in wild-type animals reduces feeding rate likely via inhibition of DAF-2/InsR signaling in PENs. In contrast, each pBoc event is almost always followed by an expulsion of gut contents in animals that express DAF-2(DN) in HENs (Figure S2D-S2E), which suggests that the communication between the intestinal defecation pacemaker and the HENs does not require active insulin signaling in the hindgut-innervating neurons. Hence, we conclude that the control of PENs and HENs by DAF-16/FoxO activation in the midgut involves independent signaling axes.

**Figure 2.**
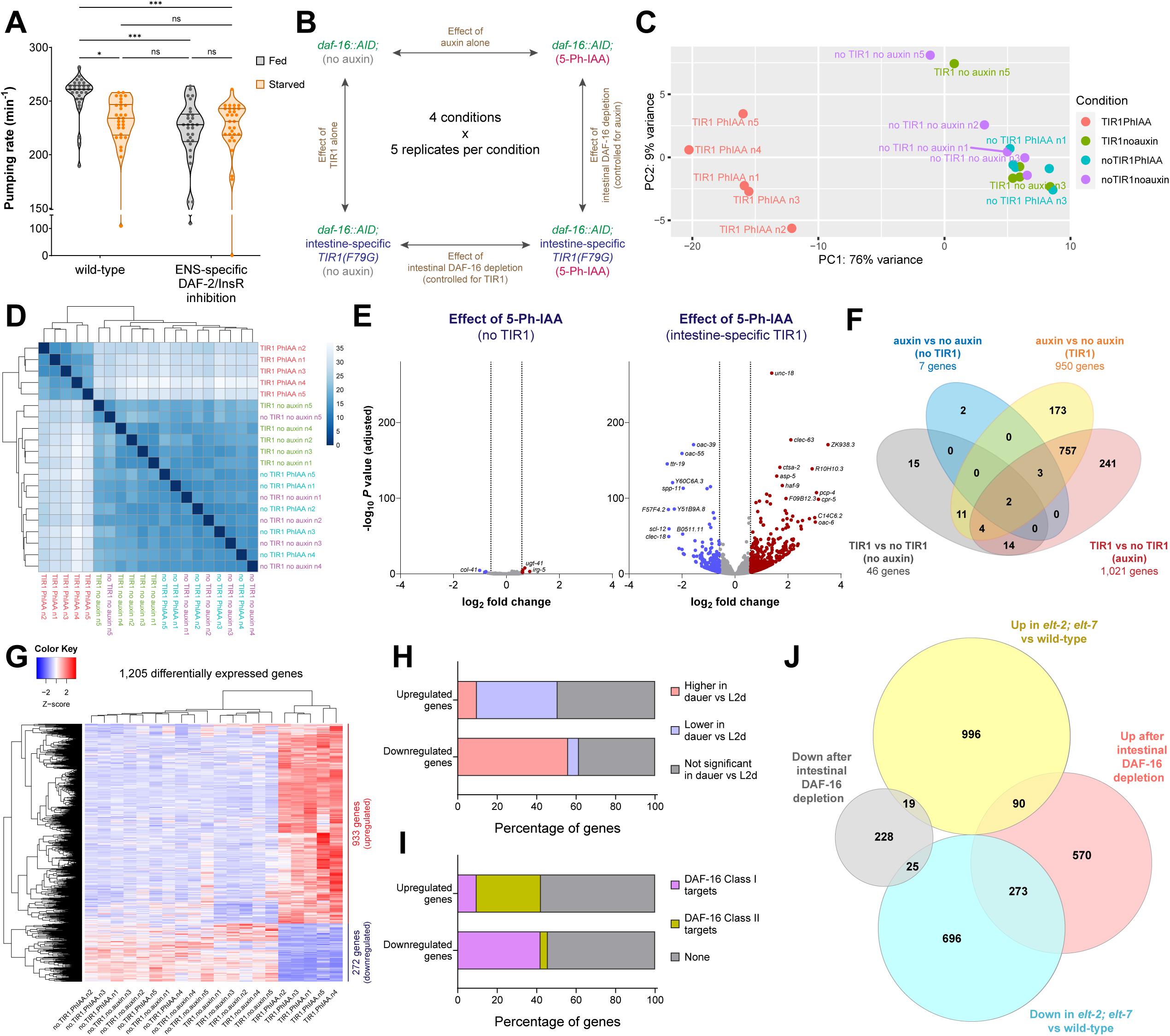
DAF-16/FoxO silences transcription of several metabolic pathway genes in the gut upon acute starvation. (A) Pharyngeal pumping rate on food in fed vs starved (4 hr) wild-type and *otIs913[pha-4prom2::daf-2(DN)]* adults. Horizontal line in the middle of data points and additional horizontal lines represent median of biological replicates, and 25^th^ and 75^th^ percentiles, respectively. *, *** and ns represent P < 0.05, P < 0.001 and not significant, respectively, in Sidak’s multiple comparisons test after two-way ANOVA. (B) Schematic of RNA-seq strategy to determine gene expression changes mediated by intestinal DAF-16 in dauer stage animals. (C) Principal component analysis (PCA) plot for all samples used in RNA-seq analysis. (D) Heatmap of sample-to-sample distance matrix for all samples used in RNA-seq analysis. Darker shades of blue indicate more similar gene expression profiles. (E) Volcano plots for gene expression changes due to auxin (5-Ph-IAA) alone or auxin in the presence of intestinal TIR1(F79G). Dashed vertical lines indicate the fold change cutoff of 1.5-fold for differential expression, log_2_(1.5) = 0.585. (F) Venn diagram for the number of differentially expressed genes for each of the four comparisons. (G) Heatmap of 1,205 genes that are differentially expressed after depletion of intestinal DAF-16 in dauers. (H) Overlap of genes regulated by intestinal DAF-16 in dauers (this study) with all genes that undergo transcriptional changes during dauer formation as previously reported ^48^. (I) Overlap of genes regulated by intestinal DAF-16 in dauers (this study) with known class I and class II transcriptional targets of DAF-16 as previously reported ^50^. (J) Venn diagram showing overlap of genes regulated by intestinal DAF-16 in dauers (this study) with all genes that are differentially expressed in *elt-2; elt-7* double mutant animals as previously reported ^51^. See also Figure S2 and Tables S1 and S2.

### Identification of DAF-16/FoxO-induced gene expression changes in the intestine during acute starvation

We sought to identify these gut-to-ENS interorgan signaling axes by profiling DAF-16/FoxO-dependent gene expression changes in the intestine during nutrient deprivation. For this comparison, we chose animals arrested in the dauer diapause stage because: (1) the dauer stage is a more robust and stable state of nutritional stress response compared to starvation during adulthood ^38^, (2) the reduction in feeding rate is much stronger in the dauer stage, i.e., a complete silencing of feeding behavior (Figure 1B), and (3) DAF-16/FoxO removal selectively from the dauer intestine using AID2 results in a strong reversal of the silenced state of the pharynx (Figures 1G, S2F and S2G). Since the AID2 system requires two components to achieve degradation of AID*-tagged DAF-16/FoxO, TIR1(F79G) and its synthetic auxin ligand 5-Ph-IAA, our experimental condition had both the AID2 components, but we also included three controls in our analysis: only TIR1(F79G) but no auxin, only auxin but no TIR1(F79G) and neither TIR1(F79G) nor auxin (Figure 2B). Since the AID2 system has not yet been extensively used for whole transcriptome level comparisons in animals, use of these controls allows to estimate the gene expression changes due to TIR1(F79G) alone or auxin alone (Figure 2B).

We find that at the whole transcriptome level, TIR1(F79G) alone or auxin alone do not produce noticeable gene expression changes (Figures 2C and 2D). All five biological replicates of the experimental condition cluster separately from all the replicates of the three control conditions (Figures 2C and 2D). The synthetic auxin ligand, 5-Ph-IAA, alters gene expression only when acting together with TIR1(F79G), but not alone (Figure 2E). On estimation of the number of differentially expressed genes for each comparison, we found that auxin alone or TIR1(F79G) alone altered the expression of less than 50 genes, while when both of the AID2 components were acting together, the expression of ∼1,000 genes was affected (Figure 2F). To find the genes that are affected by removal of DAF-16/FoxO from the intestine, we selected the 1,205 transcripts that were differentially expressed in the experimental condition in comparison to either the no TIR1(F79G) control or the no auxin control (Figures 2F and 2G). Among these 1,205 genes, 933 were upregulated and 272 were downregulated (Figure 2G and Table S1).

We compared the genes identified in our transcriptomic analysis to another study that has profiled the transcriptomes of the dauer and predauer L2D stages ^48^. In the L2D stage, the animal is uncommitted to dauer formation and can resume reproductive growth if conditions become favorable ^49^. We found that genes that are upregulated during dauer formation are significantly enriched among genes that are downregulated after DAF-16/FoxO removal from the dauer intestine (152 out of 272, *P* = 2.53 x 10^-69^) (Figure 2H). Similarly, genes that are downregulated during dauer formation are significantly enriched among genes that are upregulated after intestine-specific removal of DAF-16/FoxO (382 out of 933, *P* = 5.46 x 10^-110^) (Figure 2H), which suggests that depleting DAF-16/FoxO only from the intestine can reverse several of the gene expression changes associated with dauer formation.

The transcriptional targets of DAF-16/FoxO are classified into class I and class II ^50^. Class I targets of DAF-16 are upregulated by this transcription factor, while class II targets are downregulated ^50^. We found that genes upregulated after DAF-16/FoxO removal from the dauer intestine have a significant enrichment of class II DAF-16 targets (304 out of 933, *P* = 4.53 x 10^-96^) (Figure 2I). Similarly, genes that were downregulated after DAF-16/FoxO removal from the dauer intestine have a significant enrichment of class I DAF-16 targets (114 out of 272 = 8.86 x 10^-50^) (Figure 2I), which demonstrates that a large proportion of the genes identified in our study are direct transcriptional targets of DAF-16/FoxO.

We performed pathway enrichment analysis on genes that are upregulated after gut-specific DAF-16/FoxO removal in dauers and found that these genes are involved in several metabolic functions of the gut, such as synthesis and processing of fatty acids and vitamins, and function of organelles such as lysosomes and peroxisomes (Figure S2H). To identify if DAF-16/FoxO cooperates with other transcription factors in the gut to alter the expression of genes related to intestinal function, we scanned the upstream promoter regions of these genes for enrichment of any known transcription factor binding sites. Genes that are upregulated after DAF-16/FoxO depletion from the dauer intestine show an enrichment for the binding sites of GATA and NHR family transcription factors in their upstream sequences (Table S2). Since the GATA transcription factors ELT-2 and ELT-7 and several members of the NHR family of transcription factors are required for the expression of metabolism-related genes in intestinal epithelial cells ^51–54^, we speculated that DAF-16/FoxO shuts down several functional processes of the gut during acute nutritional deprivation by antagonizing the ability of these gut-specific transcription factors to activate specific target genes. To test this, we compared the genes that we identified to be affected by intestine-specific DAF-16/FoxO depletion with genes that are differentially expressed in animals that lack ELT-2 and ELT-7, GATA transcription factors that contribute to the differentiation program of the *C. elegans* intestine (Figure 2J)^51^. We found that approximately 30% of genes (273 out of 933, *P* = 8.23 x 10^-136^) that are upregulated after DAF-16/FoxO depletion from the dauer intestine are downregulated in animals that lack both ELT-2 and ELT-7 (Figure 2J), suggesting that DAF-16/FoxO induces global downregulation of several intestinal differentiation genes by antagonizing the activity of gut-specific GATA transcription factors during acute starvation.

For genes that are downregulated after DAF-16/FoxO depletion from the dauer intestine, their upstream sequences show the expected enrichment of the DAF-16 binding element (DBE) ‘GTAAACA’, which is similar to the binding sites of other forkhead box (FOX) family of transcription factors (Table S2) ^50,55^. In addition, we see enrichment of ‘TGCACTT’, the binding site for DAF-12/VDR, which is an NHR family transcription factor that cooperates with DAF-16/FoxO to remodel tissues during dauer formation (Table S2)^39,56^. We also found enrichment of binding sites for GATA transcription factors in the upstream sequences of genes that are downregulated after gut-specific DAF-16/FoxO depletion (Table S2). A small but statistically significant proportion of these genes (25 out of 272, *P* = 0.0045) are also downregulated in animals that lack ELT-2 and ELT-7, and many of these genes (e.g., *mtl-1*, *ftn-1*, *gst-38*, *hsp-12.3* and *hsp-12.6*) are involved in responding to stressful conditions (Figure 2J). This suggests that DAF-16/FoxO cooperates not just with another dauer-associated transcription factor DAF-12/VDR, but possibly also with GATA transcription factors such as ELT-2 to upregulate stress responsive genes in the intestine during acute nutritional deprivation, as previously shown in the context of lifespan regulation for some of the same genes (*mtl-1* and *hsp-12.6*)^57^.

### DAF-16/FoxO shuts down the expression of the gut-derived insulin peptide INS-7 to silence foregut contractions during acute starvation

Since our primary goal of the transcriptomic analysis was to identify changes in expression of secreted factors from the gut that act via the DAF-2/InsR receptor in PENs, we focused on the expression change of insulin family of genes after intestine-specific DAF-16/FoxO removal during starvation. The *C. elegans* genome encodes 39 members of the insulin family of peptides that show a complex and dynamic pattern of expression ^40,58–60^. We found that gut-specific removal of DAF-16/FoxO during acute starvation upregulates transcript levels of *ins-7* and *ins-33*, and downregulates the transcript levels of *ins-1*, *ins-18* and *ins-24* (Figure 3A). Compared to the reported changes of insulin gene expression during dauer formation ^48^, we found that they are affected in the opposite direction, i.e., *ins-1*, *ins-18* and *ins-24* are upregulated, while *ins-7* and *ins-33* are downregulated in dauers (Figure 3B), which demonstrates that the expression change of these insulins during acute starvation is mediated via DAF-16/FoxO activity in the intestine.

**Figure 3.**
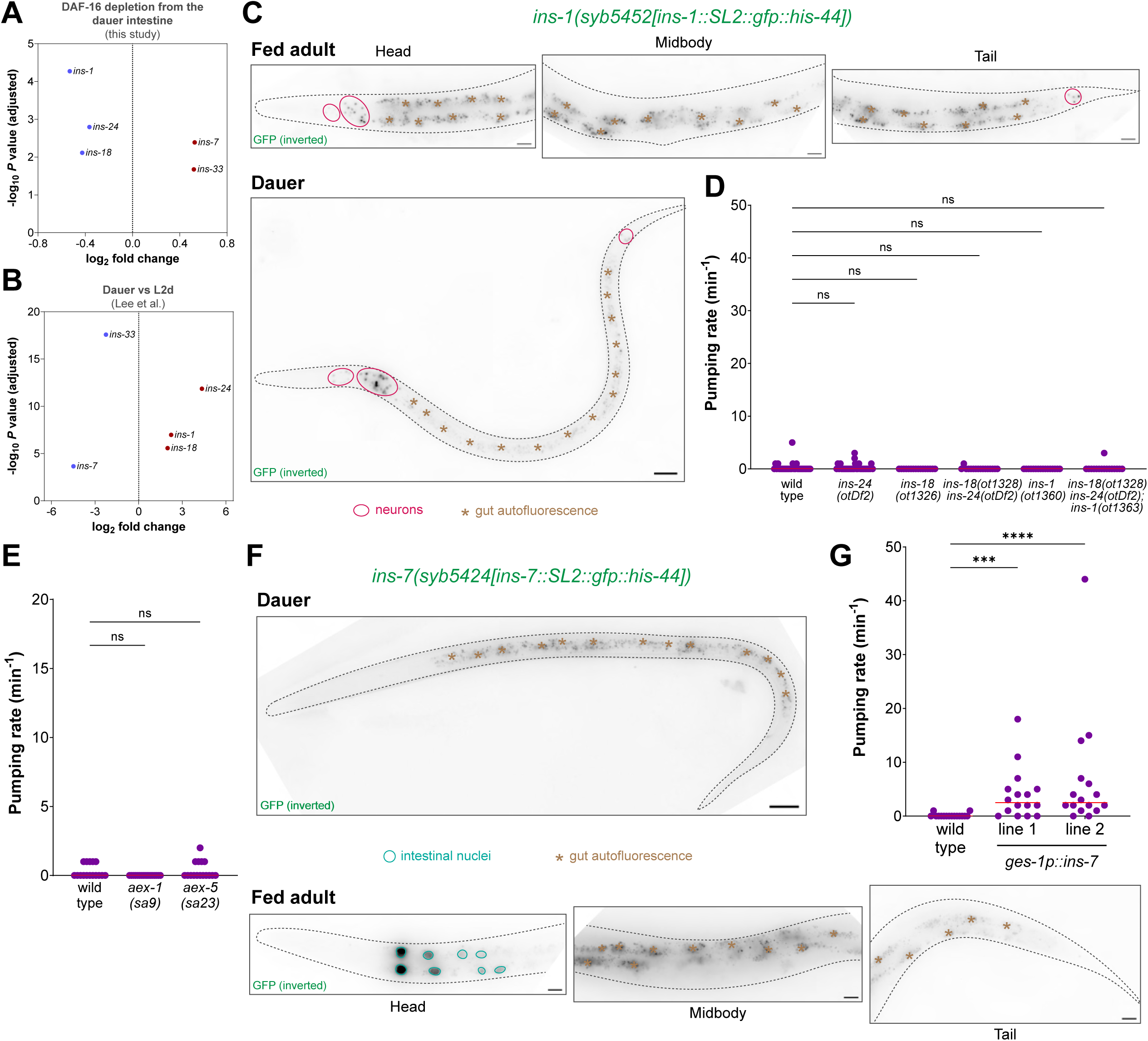
Reversing the intestinal DAF-16/FoxO-induced changes in the expression of insulin peptide genes mildly reverses the silenced state of the dauer pharynx. (A) Change in expression levels of insulin family genes that are differentially expressed after intestine-specific DAF-16 depletion in dauers. (B) Change in expression levels of insulin family genes shown in (A) from the previously reported dauer vs L2d dataset ^48^. (C) Expression of the endogenously tagged *ins-1* reporter allele *syb5452[ins-1::SL2::gfp::his-44]* in fed adult and starvation-induced dauer stage animals. Representative images of 15 animals per condition. Scale bars, 20 μm. (D) Pharyngeal pumping rate on food in starvation-induced dauer stage animals with single or combination of null mutations in *ins-1, ins-18* and *ins-24* genes. (E) Pharyngeal pumping rate on food in starvation-induced dauer stage animals with defective neuropeptide secretion from the intestine, i.e., *aex-1(sa9)* and *aex-5(sa23)* mutants. (F) Expression of the endogenously tagged *ins-7* reporter allele *syb5424[ins-7::SL2::gfp::his-44]* in fed adult and starvation-induced dauer stage animals. Representative images of 15 animals per condition. Scale bars, 20 μm. (G) Pharyngeal pumping rate on food in starvation-induced dauer stage animals that constitutively express *ins-7* in the intestine (*otEx8240* and *otEx8241* lines). Horizontal line in the middle of data points represents median value of biological replicates in (D, E, G). ***, **** and ns represent *P* < 0.001, *P* < 0.0001 and not significant, respectively, in Dunn’s multiple comparison test after Kruskal-Wallis test in (D, E, G). See also Figure S3.

Since members of the insulin peptide family can act as agonists or antagonists of DAF-2/InsR receptor to produce a combinatorial effect on insulin/IGF1 signaling at the systemic level ^34,60–62^), there are two possible scenarios for how insulin peptides made in the gut can affect ENS output: (1) agonist insulins are released from the gut in favorable conditions that act on DAF-2/InsR in the PENs to prevent DAF-16/FoxO activation and generate normal feeding behavior in the presence of abundant food, and (2) antagonist insulins are released from the gut in unfavorable conditions that inhibit DAF-2/InsR signaling in the PENs resulting in DAF-16/FoxO activation, which inhibits ENS output during periods of nutritional adversity (Figure S3A). DAF-16/FoxO activation in the intestine during starvation conditions is expected to shut down the expression of the agonist insulins and increase the expression of the antagonist insulins (Figure S3A). Since the expression of *ins-1*, *ins-18* and *ins-24* are upregulated by intestinal DAF-16 during acute starvation (Figures 3A and 3B), we speculated that active release of these insulins in the dauer stage might be required to completely silence foregut contractions. We generated CRISPR-based endogenous expression reporters for these three insulin genes, and surprisingly, none of these are expressed in the intestine in well-fed or starved conditions (Figures 3C and S3B-S3C). All of these insulins are expressed in head neurons, while *ins-1* and *ins-18* are also expressed in tail neurons (Figures 3C, and S3B-S3C), which suggests that intestinal DAF-16/FoxO activation in the gut epithelia non-cell autonomously increases the expression of *ins-1*, *ins-18* and *ins-24* in the nervous system during acute starvation. We generated CRISPR-based null alleles for all of these insulin genes and found that deletion of *ins-1*, *ins-18* and *ins-24* either individually or in combination does not reverse the silenced state of the dauer pharynx (Figure 3D), suggesting that upregulation of these insulin peptides is not required to silence foregut contractions during acute starvation.

We found that a large number of non-insulin neuropeptide genes are transcriptionally downregulated when DAF-16/FoxO is depleted from the dauer intestine (Figure S3D). This is consistent with a previous study that showed that the entire family of FMRFamide-like neuropeptide (*flp*) genes are transcriptionally upregulated during dauer formation ^48^, which suggests that some of these neuropeptides might be involved in silencing ENS output during starvation. We tested whether blocking pathways related to neuropeptide processing or release affects the silenced state of the dauer pharynx. We found that disruption of *unc-31/CAPS*, a calcium-dependent activator protein required for the release of neuropeptides in dense-core vesicles ^63,64^, *sbt-1/SCG5*, a chaperone for proprotein convertases that cleave propeptides ^65^, or *trap-1/SSR1*, an ER membrane protein required for biogenesis of antagonistic insulin peptides ^66^, did not induce foregut contractions in dauer stage animals (Figure S3E). Since *unc-31/CAPS*, *sbt-1/SCG5* and *trap-1/SSR1* are involved in the processing and release of neuropeptides from the nervous system but the signal that silences ENS output is regulated by DAF-16/FoxO in the gut, we also manipulated the genes that are involved in secretion of neuropeptides from the intestinal epithelial cells. Disrupting the functions of *aex-1/UNC13D* or *aex-5/PCSK5*, constituents of the neuropeptide secretion machinery of the gut ^20,67^, did not generate contractions in the silenced dauer pharynx (Figure 3E), which indicates that active secretion of neuropeptides from the gut epithelial cells is likely not required to silence PEN activity during acute starvation.

Next, we focused on *ins-7* and *ins-33*, insulin genes that are downregulated by intestinal DAF-16/FoxO in dauer stage animals (Figures 3A and 3B). We generated CRISPR-based endogenous expression reporters for both these genes and found that *ins-7* is only expressed in the anterior region of the midgut of well-fed adults, while *ins-33* is expressed in hypodermal cells, but not in the gut (Figures 3F and S3F). During acute starvation, expression of both *ins-7* and *ins-33* are reduced in the respective tissues (Figures 3F and S3F). We found that constitutive expression of *ins-7* from gut epithelia produces a mild but significant reversal of the silenced dauer pharynx, while constitutive expression of *ins-33* produces no effect (Figures 3G and S3G). This indicates that release of the INS-7 peptide from gut epithelia can induce foregut contractions in starvation conditions that completely silence ENS output.

### DAF-16/FoxO shuts down the neuropeptide secretion machinery of the gut during acute starvation

In addition to altering the expression of neuropeptides, we found that depletion of DAF-16/FoxO from the dauer intestine also upregulates the expression of several genes that encode for components of the neuropeptide secretion machinery of the gut (Figure 4A). Among these genes, the strongest upregulation is observed for *aex-1/UNC13D*, a SNARE regulator ^20,67^, while modest upregulation is found for *aex-4/SNAP25*, a t-SNARE protein ^20^, and *aex-5/PCSK5*, a proprotein convertase (Figure 4A)^20,67^. All of these *aex* genes are significantly downregulated during dauer formation (Figure 4B)^48^. We generated CRISPR-based endogenous reporters for these *aex* genes to profile their expression pattern in well-fed adults, starvation-induced dauers and during the L3 larval stage, which is the developmental equivalent of the dauer stage in the reproductive phase of growth. For *aex-1/UNC13D*, we observe strong expression in all intestinal epithelial cells in adults and L3 larvae grown in nutrient replete conditions (Figure 4C). In well-fed adults, we also observe *aex-1/UNC13D* expression in uterine cells and dim expression in the hypodermis (Figure 4C). In dauers, we observe a strong reduction in *aex-1/UNC13D* expression in the intestinal epithelium (Figures 4C and 4D). For both *aex-4/SNAP25* and *aex-5/PCSK5*, we observe strong expression in the epithelial cells of the midgut during adulthood and L3 larval stage (Figures 4E and S4A). In well-fed adults, *aex-4/SNAP25* is also expressed in uterine cells, while *aex-5/PCSK5* is also expressed in the body wall muscles of animals (Figures 4E and S4A). In the dauer stage, both *aex-4/SNAP25* and *aex-5/PCSK5* undergo a strong reduction in expression in the intestinal epithelial cells (Figures 4E and 4F, and S4A and S4B). In summary, DAF-16/FoxO strongly downregulates the expression of these *aex* genes in the gut epithelia during acute starvation (Figures 4A-4F, and S4A and S4B).

**Figure 4.**
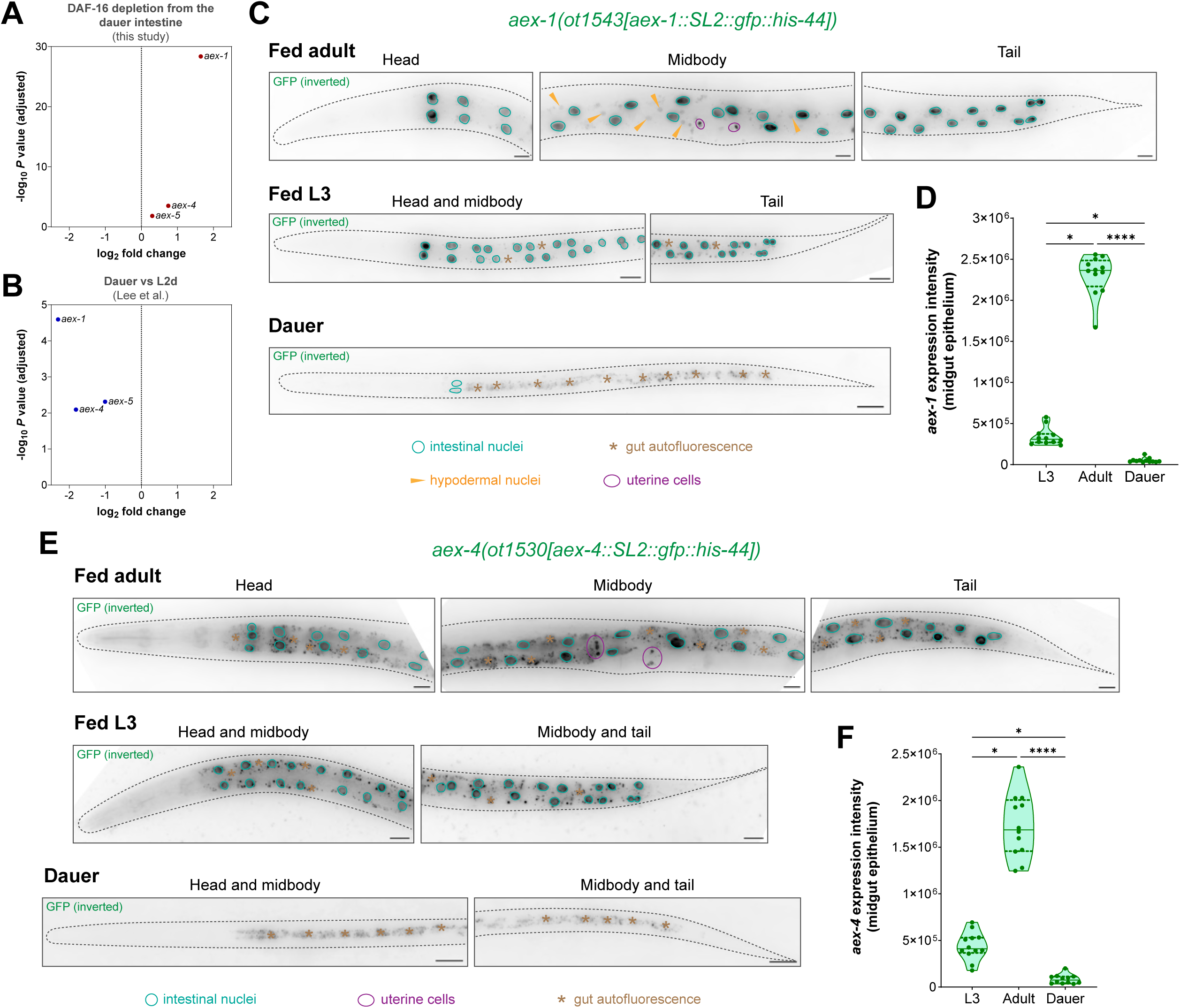
DAF-16/FoxO activation shuts down the intestinal neuropeptide secretion machinery genes during prolonged starvation. (A) Change in expression levels of *aex-1*, *aex-4* and *aex-5* genes after intestine-specific DAF-16 depletion in dauers. (H) Change in expression levels of *aex-1*, *aex-4* and *aex-5* genes from the previously reported dauer vs L2d dataset ^48^. (A) Expression of the endogenously tagged *aex-1* reporter allele *aex-1(ot1543[aex-1::SL2::gfp::his-44])* in fed adult and L3 larva, and in starvation-induced dauer stage animals. Scale bars, 20 μm. (A) Quantification of *aex-1* expression in midgut epithelial cells of fed adult and L3 larva, and starvation-induced dauer stage animals. (A) Expression of the endogenously tagged *aex-4* reporter allele *aex-4(ot1530[aex-4::SL2::gfp::his-44])* in fed adult and L3 larva, and in starvation-induced dauer stage animals. Scale bars, 20 μm. (A) Quantification of *aex-4* expression in midgut epithelial cells of fed adult and L3 larva, and starvation-induced dauer stage animals. Horizontal line in the middle of data points and additional horizontal lines represent median of biological replicates, and 25^th^ and 75^th^ percentiles, respectively in (D, F). * and **** represent *P* < 0.05 and *P* < 0.0001, respectively, in Dunn’s multiple comparison test after Kruskal-Wallis test in (D, F). See also Figure S4.

From existing ChIP-seq datasets, we found that the upstream genomic regions of *aex-1/4/5* genes show DAF-16/FoxO binding peaks (Figures S4C-E). These regions also show binding peaks for ELT-2, a GATA transcription factor that is the master regulator of intestinal differentiation (Figures S4C-E)^52^. This suggests that ELT-2 might be required for the expression of *aex-1/4/5* genes for normal secretory functions of the intestinal epithelia in favorable conditions, while DAF-16/FoxO activation antagonizes the activity of this gut-specific GATA transcription factor to silence the expression of *aex-1/4/5* genes during nutritional adversity.

Since multiple components of the neuropeptide secretion machinery of the intestine are transcriptionally silenced during starvation, we generated a secretion reporter to directly test whether the release of neuropeptides from gut epithelia is hindered in adverse conditions. We generated a transgene with the insulin peptide INS-1 translationally fused to a TagRFP fluorescent protein and expressed it specifically in intestinal epithelial cells (Figure 5A). From the same construct, a GFP-tagged histone H2B protein is synthesized after an SL2 trans-splicing event, which reports whether transcription of the transgene is affected in any condition (Figure 5A). In well-fed adults and L3 larvae, the INS-1::TagRFP peptide is synthesized only in gut epithelia, but most of this protein is secreted from the basolateral side of these cells and the tagged INS-1 peptide accumulates in the coelomocytes, macrophage-like scavenger cells in the body coelom (Figure 5A). In the dauer diapause stage, we observe that instead of being released from the basolateral side of gut epithelia, the INS-1::TagRFP peptide is now released into the lumen of the gut from the apical side of intestinal cells (Figures 5A and 5B), a phenomenon that has been reported previously but the physiological relevance was unknown ^68^. In addition, we observe strong accumulation of the tagged INS-1 peptide in the cytoplasm of gut epithelial cells in dauers (Figures 5A and 5B), suggesting that the secretion of this neuropeptide is severely reduced during acute starvation conditions. We tested whether this neuropeptide secretion block in the dauer intestine is a direct consequence of DAF-16/FoxO activation. After gut-specific depletion of DAF-16/FoxO in dauers, we observe a robust increase in the secretion of INS-1::TagRFP from the basolateral side of intestinal cells with reliably stronger accumulation in the coelomocytes of the animal (Figures 5C and 5D). These findings suggest that DAF-16/FoxO cell autonomously silences the secretion of neuropeptides from the intestine into the body coelom during acute starvation.

**Figure 5.**
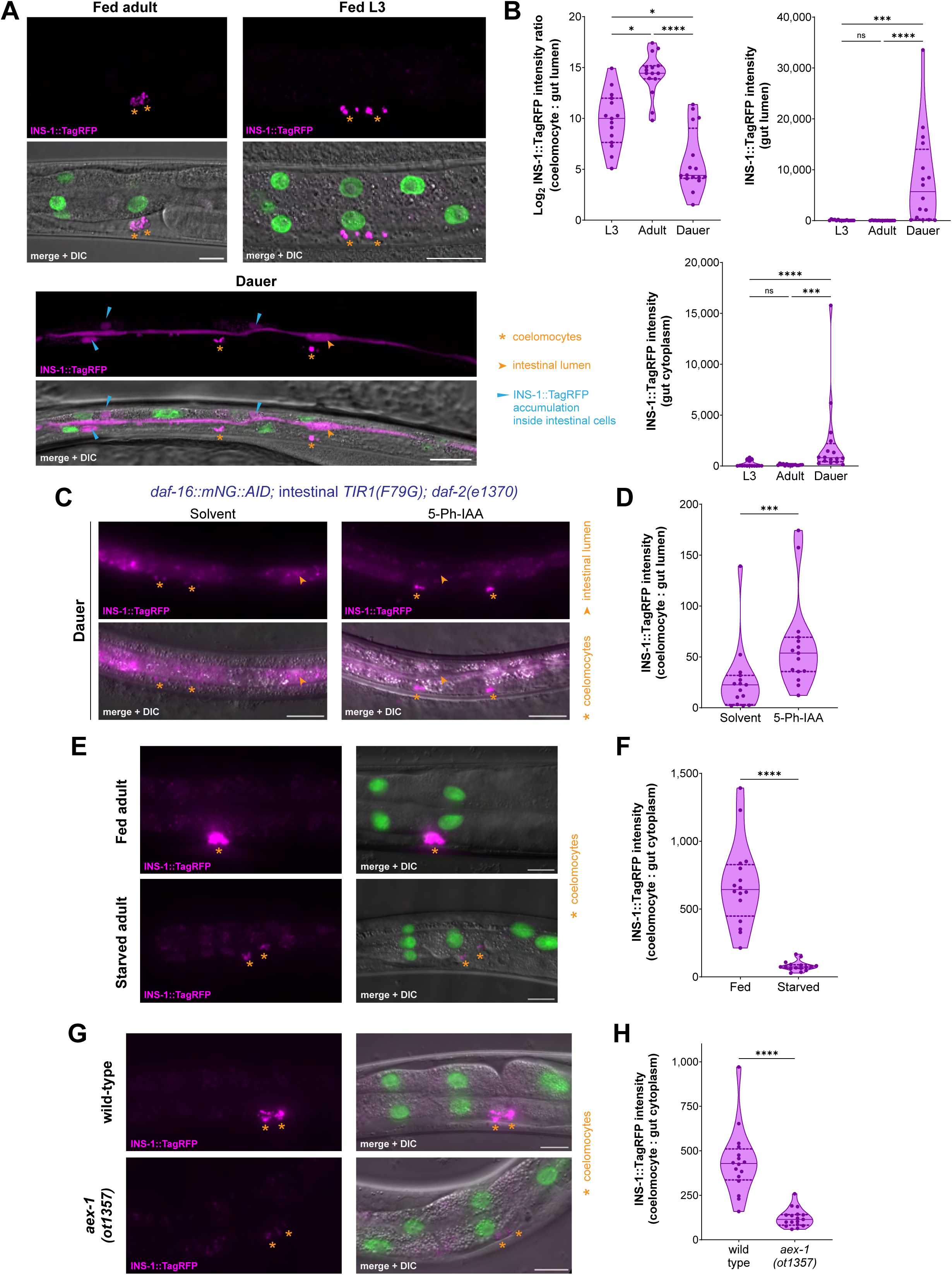
DAF-16/FoxO shuts down the release of insulin peptides from the midgut epithelium during prolonged starvation. (A) Secretion of INS-1::TagRFP from the intestine into coelomocytes in fed adult and L3 larva, and in starvation-induced dauer stage animals expressing *otEx8059[ges-1p::ins-1::tagRFP::SL2::gfp::his-44]*. Scale bars, 20 μm. (B) Quantification of INS-1::TagRFP intensity in gut lumen and gut cytoplasm, and the ratio of INS-1::TagRFP intensity in coelomocytes: gut cytoplasm in fed adult and L3 larva, and in starvation-induced dauer stage animals expressing *otEx8059[ges-1p::ins-1::tagRFP::SL2::gfp::his-44]*. (C) Secretion of INS-1::TagRFP from the intestine into coelomocytes in *daf-16(ot853); otSi2[ges-1p::TIR1(F79G)]; daf-2(e1370); otEx8220[ges-1p::ins-1::tagRFP::SL2::ebfp2::his-44]* dauer stage animals after intestine-specific DAF-16 depletion. Animals were treated with either solvent (ethanol) or 100 μM 5-Ph-IAA. Scale bars, 20 μm. (D) Ratio of INS-1::TagRFP intensity in coelomocytes: gut lumen in *daf-16(ot853); otSi2[ges-1p::TIR1(F79G)]; daf-2(e1370); otEx8220[ges-1p::ins-1::tagRFP::SL2::ebfp2::his-44]* dauer stage animals after intestine-specific DAF-16 depletion. Animals were treated with either solvent (ethanol) or 100 μM 5-Ph-IAA. (E) Secretion of INS-1::TagRFP from the intestine into coelomocytes in fed vs starved (24 hr) adults expressing *otIs904[ges-1p::ins-1::tagRFP::SL2::gfp::his-44]*. Scale bars, 20 μm. (F) Ratio of INS-1::TagRFP intensity in coelomocytes: gut cytoplasm in fed vs starved (24 hr) adults expressing *otIs904[ges-1p::ins-1::tagRFP::SL2::gfp::his-44]*. (G) Secretion of INS-1::TagRFP from the intestine into coelomocytes in fed wild-type or *aex-1(ot1357)* adults expressing *otIs904[ges-1p::ins-1::tagRFP::SL2::gfp::his-44]*. Scale bars, 20 μm. (H) Ratio of INS-1::TagRFP intensity in coelomocytes: gut cytoplasm in fed wild-type or *aex-1(ot1357)* adults expressing *otIs904[ges-1p::ins-1::tagRFP::SL2::gfp::his-44]*. Horizontal line in the middle of data points and additional horizontal lines represent median of biological replicates, and 25^th^ and 75^th^ percentiles, respectively in (B, D, F, H). *, ***, **** and ns represent *P* < 0.05, *P* < 0.001, *P* < 0.0001 and not significant, respectively, in Dunn’s multiple comparison test after Kruskal-Wallis test in (B) and in Mann-Whitney test in (D, F, H). See also Figure S5.

Next, we tested whether neuropeptide secretion from the gut epithelial cells is blocked only during the dauer diapause state or also when adult animals are subjected to acute starvation. We found that extended food deprivation in adults strongly reduces the basolateral secretion of the tagged INS-1 peptide from the gut, which is apparent from a much reduced INS-1::TagRFP accumulation in the coelomocytes (Figures 5E and 5F). The lower insulin signal in the coelomocytes of starved adults is not due to reduced expression of this construct in the gut because the intensity of GFP-tagged H2B in the intestinal nuclei remains unchanged during starvation (Figure S5A). Since *aex-1/UNC13D* shows a strong DAF-16/FoxO-induced silencing during dauer formation (Figures 4A-4D), which is associated with a neuropeptide secretion block from the dauer intestine (Figures 5A-5D), we asked whether *aex-1/UNC13D* depletion also inhibits neuropeptide release from the gut of adults. We generated a CRISPR-based deletion of the entire *aex-1/UNC13D* protein coding region and found much reduced secretion of the tagged INS-1 peptide from the intestinal epithelial cells into the coelomocytes in *aex-1/UNC13D* null animals even in nutrient rich conditions (Figures 5G and 5H). These results collectively demonstrate a role of *aex-1/UNC13D* in neuropeptide release from gut epithelia in well-fed conditions, and DAF-16/FoxO mediated silencing of *aex-1/UNC13D* strongly reduces the secretion of neuropeptides from the gut during acute starvation conditions.

### Active secretion of INS-7 and INS-35 from gut epithelia is essential for normal foregut contractility in favorable conditions

Since starvation-induced DAF-16/FoxO activity in the gut reduces intestinal neuropeptide secretion as well as the output from PENs in terms of foregut contractions, we speculated that active neuropeptide release from the gut is required for generating normal feeding behavior in favorable conditions. We found that animals with previously described loss-of-function mutations in the intestinal secretion machinery genes ^20,67^, *aex-1/UNC13D* and *aex-5/PCSK5*, have a significantly reduced rate of foregut contractions compared to wild-type even in nutrient replete conditions (Figure 6A). This phenotype was also observed in animals with the CRISPR-generated null allele of *aex-1/UNC13D* (Figure 6B). Since *aex-1/UNC13D* and *aex-5/PCSK5* mutants were initially identified in a genetic screen for animals with defective defecation behavior ^22^, it can be postulated that these animals are feeding less because of their lower rate of expulsion from the hindgut. Though previous studies have observed no correlation between the rates of foregut and hindgut contractions ^69^, we tested whether a defecation defective strain that does not affect neuropeptide secretion from the midgut can affect the rate of feeding. For this assay, we used a loss-of-function allele of *nlp-40*, a neuropeptide that is rhythmically released from the intestinal epithelia to activate the HENs and initiate expulsion of gut contents ^19^. Unlike the *aex-1/UNC13D* and *aex-5/PCSK5* mutants, depletion of *nlp-40* does not affect the release of other neuropeptides from the intestinal cells, even though all of these strains have strong defects in expulsion behavior ^19,20^. We found that animals lacking NLP-40 show the same frequency of foregut contractions as wild-type animals (Figure S5B), indicating that inhibiting defecation behavior without disrupting the neuropeptide secretion machinery of the gut does not affect the output from PENs.

**Figure 6.**
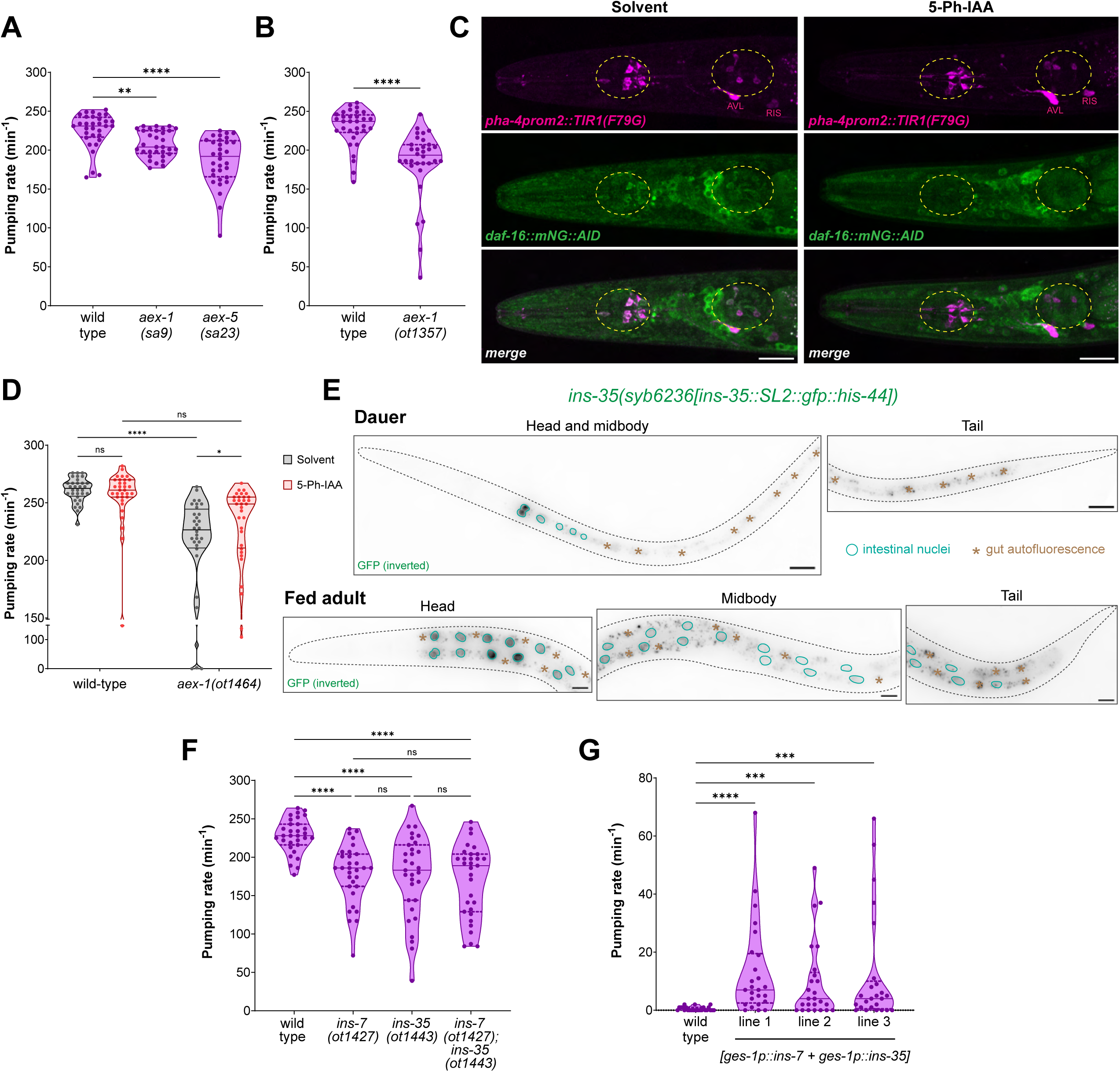
Active intestinal secretion of INS-7 and INS-35 is essential for normal pharyngeal pumping behavior in favorable conditions. (A) Pharyngeal pumping rate on food in fed wild-type, *aex-1(sa9)* and *aex-5(sa23)* adults. (B) Pharyngeal pumping rate on food in fed wild-type and *aex-1(ot1357)* adults. (C) DAF-16 depletion from enteric neurons of the foregut in *daf-16(ot853); otIs908[pha-4prom2::TIR1(F79G)]* adults. Animals were treated with either solvent (ethanol) or 100 μM 5-Ph-IAA. DAF-16 was not detected in enteric neurons of the foregut after 5-Ph-IAA treatment in 15/15 animals. The anterior and posterior bulbs of the pharynx are shown with dashed circles. Non-pharyngeal neurons where *otIs908* is expressed are labeled. Scale bars, 20 μm. (D) Pharyngeal pumping rate on food after DAF-16 depletion from enteric neurons of the foregut in fed *daf-16(ot853); otIs908[pha-4prom2::TIR1(F79G)]* adults in wild-type or *aex-1(ot1464)* genetic background. Animals were treated with either solvent (ethanol) or 100 μM 5-Ph-IAA. (E) Expression of the endogenously tagged *ins-35* reporter allele *syb6236[ins-35::SL2::gfp::his-44]* in fed adult and starvation-induced dauer stage animals. Representative images of 15 animals per condition. Scale bars, 20 μm. (F) Pharyngeal pumping rate on food in fed wild-type, *ins-7(ot1427)*, *ins-35(ot1443)* and *ins-7(ot1427); ins-35(ot1443)* adults. (G) Pharyngeal pumping rate on food in starvation-induced dauer stage animals that constitutively co-express *ins-7* and *ins-35* in the intestine (*otEx8222*, *otEx8223* and *otEx8224*). Horizontal line in the middle of data points and additional horizontal lines represent median of biological replicates, and 25^th^ and 75^th^ percentiles, respectively in (A, B, D, F, G). *, **, ***, **** and ns represent *P* < 0.05, *P* < 0.01, *P* < 0.001, *P* < 0.0001 and not significant, respectively, in Dunn’s multiple comparison test after Kruskal-Wallis test in (A, F, G), in Mann-Whitney test in (B) and in Sidak’s multiple comparisons test after two-way ANOVA in (D). See also Figure S5.

Since the intestine-to-pharynx interorgan signal is likely an insulin peptide that acts via the DAF-2/InsR receptor to maintain DAF-16/FoxO in an inactive state in PENs in order to generate normal foregut contractions in nutrient-rich conditions (Figures 2A and S2B), we hypothesized that removing DAF-16/FoxO specifically from the ENS should reverse the reduced feeding rate in animals that have defective neuropeptide secretion from the midgut. To validate this, we expressed *TIR1(F79G)* specifically in the ENS of an animal that has its endogenous *daf-16/FoxO* locus tagged with *AID\** (Figure 6C). In the presence of the auxin ligand, AID*-tagged endogenous DAF-16/FoxO protein is specifically depleted from the ENS of the animal (Figure 6C). We found that ENS-specific DAF-16/FoxO depletion does not affect the rate of feeding in wild-type animals, but it significantly increases the rate of foregut contractions in the intestinal neuropeptide secretion defective *aex-1/UNC13D* null animals (Figure 6D). After depleting DAF-16/FoxO specifically from the ENS of *aex-1/UNC13D* null animals, their feeding rates become similar to that of wild-type animals grown in the abundance of food (Figure 6D). This increase in the rate of foregut contractions is not observed in *aex-1/UNC13D* null animals that are treated with the same dose of auxin in the absence of TIR1(F79G) (Figure S5C), ruling out the possibility that suppression of this phenotype is due to treatment with the synthetic auxin ligand. This suggests that a feeding defect due to the absence of neuropeptide secretion from midgut epithelial cells can be completely suppressed by removal of a transcription factor from the PENs, indicating the presence of a midgut-to-ENS communication that is modulated during nutritional adversity.

To identify the molecular nature of the interorgan signal from the gut, we interrogated a recent single-cell RNA-seq dataset that captured gene expression from all major cell types in *C. elegans* ^70^. We found that only two insulin genes, *ins-7* and *ins-35*, are specifically enriched in the intestinal cluster (Figure S5D). We confirmed using a CRISPR-based endogenous expression reporter that *ins-7* is only expressed in the anterior region of the midgut of well-fed adults and its expression is strongly reduced during acute starvation (Figure 3F). We also generated a CRISPR-based reporter for *ins-35* and found that this insulin peptide is only expressed in all of the midgut epithelial cells of the animal in well-fed conditions, and in no other tissue type (Figure 6E). In the dauer diapause stage, *ins-35* expression is restricted to the anterior cells of intestinal epithelia (Figure 6E), but at the functional level, INS-35 secretion from the basolateral side will be much reduced due to the neuropeptide secretion block in acute starvation conditions (Figures 5A and 5B). To determine whether these insulin peptides synthesized in the intestine constitute the gut-to-pharynx interorgan signal, we generated CRISPR-based null alleles for both *ins-7* and *ins-35*. We found that the rate of foregut contractions in nutrient replete conditions is significantly reduced in the absence of either INS-7 or INS-35, and if both insulin peptides are removed simultaneously, we do not observe an additive effect (Figure 6F). These results indicate that INS-7 and INS-35 released from the gut epithelial cells are required for normal feeding behavior in favorable conditions.

We found that similar to *ins-7* (Figure 3G), constitutive expression of *ins-35* in midgut epithelial cells results in a mild but significant reversal of the silenced state of the dauer pharynx (Figure S5E). However, constitutive expression of *ins-7* and *ins-35* simultaneously in the intestine of dauers generates a much stronger output from the PENs, and we observe higher than 20 contractions per minute in the top quartile of these animals, which are still arrested in the dauer diapause stage (Figure 6G). Since the feeding behavior was recorded five minutes after these dauer stage animals were transferred to food, it could be argued that ectopic expression of *ins-7* and *ins-35* triggers a faster exit from the dauer stage when these animals encounter food. We consider this unlikely because: (1) these insulin peptides do not affect the dauer exit process ^61^, and (2) molecular and behavioral changes associated with dauer exit are observed at least an hour after the dauer stage animals encounter food ^71,72^. To rule out the possibility of a faster dauer exit in animals co-expressing *ins-7* and *ins-35* from the gut, we measured their feeding behavior in the *daf-7(e1372)* dauer-constitutive genetic background. In the presence of the *daf-7(e1372)* mutation, animals constitutively arrest in the dauer stage at 25°C independent of the presence of food ^73^. We found that in *daf-7(e1372)* dauers, constitutive co-expression of *ins-7* and *ins-35* from the intestine is sufficient to induce rapid foregut contractions (Figure S5F). These findings indicate that the gut-derived insulin peptides, INS-7 and INS-35, are not only required for normal feeding behavior in favorable environments, but also are sufficient to induce foregut contractions in harsh environmental conditions that completely silence output from the PENs.

### The serotonergic NSM neurons sense insulin signals from the gut to generate normal feeding behavior

We sought to identify the site of action of the gut-derived insulin peptides in the pharyngeal ENS. Previous studies have found that the bilaterally symmetric MC neuron pair is the pacemaker of the pharynx because their ablation strongly reduces the rate of foregut contractions ^23,25^. To test whether pacemaker activity of the MC neurons requires active DAF-2/InsR signaling, we generated a strain that expresses DAF-2(DN) only in the MC neurons in the entire ENS (Figure 7A). Inhibition of DAF-2/InsR in MC neurons did not affect the rate of foregut contractions (Figure 7B), suggesting that the MC neuron pair does not require insulin signals to generate normal feeding output in favorable conditions.

**Figure 7.**
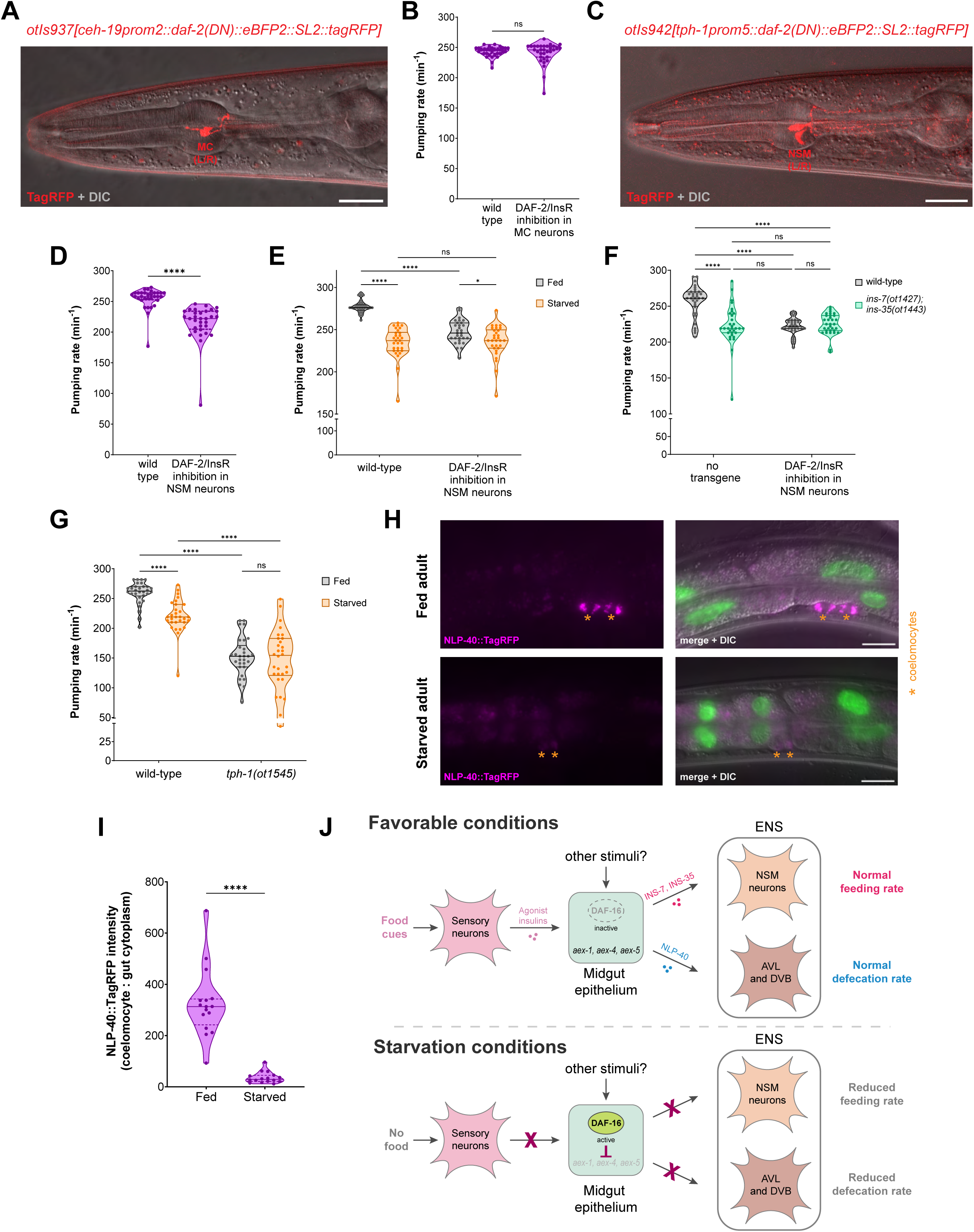
NSM senses INS-7 and INS-35 from the gut to enhance pharyngeal nervous system output during the fed state. (A) Expression of *otIs937[ceh-19prom2::daf-2(DN)::eBFP2::SL2::tagRFP]* in MC(L/R) neurons in the ENS of the foregut. Scale bar, 20 μm. (B) Pharyngeal pumping rate on food in fed wild-type and *otIs937[ceh-19prom2::daf-2(DN)::eBFP2::SL2::tagRFP]* adults. (C) Expression of *otIs942[tph-1prom5::daf-2(DN)::eBFP2::SL2::tagRFP]* in NSM(L/R) neurons in the ENS of the foregut. Scale bar, 20 μm. (D) Pharyngeal pumping rate on food in fed wild-type and *otIs942[tph-1prom5::daf-2(DN)::eBFP2::SL2::tagRFP]* adults. (E) Pharyngeal pumping rate on food in fed vs starved (4 hr) wild-type and *otIs942[tph-1prom5::daf-2(DN)::eBFP2::SL2::tagRFP]* adults. (F) Pharyngeal pumping rate on food in fed *otIs942[tph-1prom5::daf-2(DN)::eBFP2::SL2::tagRFP]* expressing adults in wild-type or *ins-7(ot1427); ins-35(ot1443)* genetic background. (G) Pharyngeal pumping rate on food in fed vs starved (4 hr) wild-type and *tph-1(ot1545)* adults. (H) Secretion of NLP-40::TagRFP from the intestine into coelomocytes in fed vs starved (8 hr) adults expressing *otIs927[ges-1p::nlp-40::tagRFP::SL2::gfp::his-44]*. Scale bars, 20 μm. (I) Ratio of NLP-40::TagRFP intensity in coelomocytes: gut cytoplasm in fed vs starved (8 hr) adults expressing *otIs927[ges-1p::nlp-40::tagRFP::SL2::gfp::his-44]*. (J) Model for regulation of both feeding and defecation behavior by neuropeptides released from midgut epithelial cells in favorable or starved conditions. Horizontal line in the middle of data points and additional horizontal lines represent median of biological replicates, and 25^th^ and 75^th^ percentiles, respectively in (B, D-G, I). *, **** and ns represent *P* < 0.05, *P* < 0.0001 and not significant, respectively, in Mann-Whitney test in (B, D, I) and in Sidak’s multiple comparisons test after two-way ANOVA in (E-G).

Next, we targeted the serotonergic NSM neuron pair in the ENS, which has a known neuromodulatory role in increasing the rate of feeding in the presence of food ^74^. To inhibit DAF-2/InsR signaling in the NSM neurons, we expressed DAF-2(DN) using a *cis*-regulatory element of the *tph-1/TPH* gene that drives expression exclusively in the NSM neurons in the entire nervous system (Figure 7C)^75^. We found that inhibition of DAF-2/InsR signaling only in the NSM neurons is sufficient to reduce the rate of feeding in nutrient replete conditions (Figure 7D).

NSM-specific DAF-2(DN) expression also strongly suppresses the effect of starvation on the rate of feeding (Figure 7E), suggesting a role of insulin signaling in NSM neurons to mediate an internal state-dependent change in feeding behavior. Compared to wild-type animals, the reduction in feeding rate due to NSM-specific DAF-2/InsR inhibition is similar to that in animals that lack INS-7 and INS-35 (Figure 7F). To confirm that the gut derived insulins, INS-7 and INS-35, act via DAF-2/InsR in NSM to regulate the rate of foregut contractions in favorable environments, we performed an epistasis experiment. We found that inhibiting DAF-2/InsR in the NSM neurons of animals that lack INS-7 and INS-35 results in no further decrease in their rate of feeding in nutrient rich conditions (Figure 7F). These results indicate that the neuromodulatory NSM neurons in the ENS sense the insulin peptides released from the midgut epithelial cells to generate rapid foregut contractions when food is abundant.

Since the NSM neurons increase the rate of foregut contractions in a serotonin (5-HT)-dependent manner ^76^, we asked whether 5-HT plays a role in altering the rate of feeding in starved animals. Previous studies have shown that animals lacking *tph-1/TPH*, the tryptophan hydroxylase enzyme essential for neuronal 5-HT biosynthesis, are unable to increase foregut contractions when they encounter food ^77^. Using a CRISPR-generated null allele of *tph-1/TPH*, we show that animals lacking the ability to synthesize 5-HT in neurons have a much lower rate of feeding than wild-type animals (Figure 7G). Unlike wild-type animals, starvation does not reduce the rate of foregut contractions in *tph-1/TPH* null animals (Figure 7G), which supports the role of serotonergic signaling in modulating feeding behavior in the context of nutritional status of the animal.

### Intestinal DAF-16/FoxO controls the second neuropeptidergic signaling axis to HENs

Previous analysis has shown that the intestinally expressed AEX-1/4/5 peptidergic release machinery acts in the same pathway as NLP-40 to activate AEX-2, the NLP-40 receptor in HENs, to control defecation behavior ^19,20^. The DAF-16/FoxO-dependent downregulation of *aex-1/4/5* under starvation conditions, described above, would therefore be expected to affect NLP-40 release from intestinal cells. To test this prediction, we generated a strain that expresses the NLP-40 neuropeptide fused with TagRFP specifically in the midgut epithelial cells. In well-fed adults, NLP-40::TagRFP synthesized in the intestine is secreted from the basolateral side of these cells into the body coelom, which is visualized by the accumulation of this neuropeptide in the coelomocytes (Figure 7H). Secretion of NLP-40 from the midgut epithelium is indeed strongly reduced after prolonged starvation in adults (Figures 7H and 7I). Our findings collectively show that DAF-16/FoxO shuts down the release of both insulin and non-insulin peptidergic signals from the midgut epithelial cells to enteric neurons in the foregut and hindgut circuits during periods of nutrient scarcity.

## DISCUSSION

How the internal state of an animal affects CNS functions has received extensive attention ^31,78^, but how internal states affect the enteric nervous system remains much less studied. Here we have shown that the epithelial cells in the *C. elegans* midgut coordinate the function of two branches of the enteric nervous system by controlling the release of distinct peptidergic signals (Figure 7J). These two distinct signaling axes – insulins from the midgut to the PENs and NLP-40 from the midgut to HENs – are coordinated via DAF-16/FoxO transcriptionally regulating key nodes of the peptidergic secretion machinery (Figure 7J). Based on available ChIP-Seq data for DAF-16/FoxO and the master regulator of intestinal differentiation, ELT-2 (Figure S4C-E), we propose that ELT-2 normally activates *aex-1/4/5* and that this activation is antagonized by DAF-16/FoxO under food-deprived conditions.

On sensing food-related cues in the environment, *C. elegans* chemosensory neurons release insulin peptides to modulate DAF-16/FoxO activity in the intestine ^34^, which, as we have shown here, regulates the release of other insulins to control pharyngeal pumping and of a non-insulin neuropeptide to control enteric muscle contractions. We hypothesize that the relay of insulin signals from sensory neurons to midgut back to enteric neurons permits midgut epithelia cells to serve as an integrator of various types of nutritional signals both from the external environment and from internal nutritional signals from digested food within the alimentary tract. The nutrient sensing TORC1 and TORC2 pathways have been shown to regulate DAF-16/FoxO activity specifically in the intestine ^79–81^, and the AAK-2/AMPK kinase directly phosphorylates DAF-16/FoxO to increase its activity during periods of extended starvation ^82^. Reduced TORC2 signaling in the gut in adverse environments inhibits the expression of insulin peptides in chemosensory neurons, which suggests a role of the intestine as an amplifier of a starvation signal since it can further reduce the release of agonist insulin peptides from the nervous system via a feed-forward loop ^83^. Hence, intestinal DAF-16/FoxO may integrate signals from multiple stress response pathways to modify nervous system function during periods of extended nutritional adversity.

It has been well documented that peptides of the insulin family can cross the blood-brain barrier to affect CNS function in mammals ^84^. Research in invertebrate models has shed light on the mechanistic bases by which the release of specific insulin peptides is actively regulated from non-neuronal tissues in the gut to modulate behaviors during different internal states of the animal. Neuropeptides released from intestinal epithelia have been shown to affect sensory behaviors such as thermotaxis and chemotaxis during prolonged starvation ^36,85^. Altered insulin signaling from the gut during periods of food scarcity can also modify secretion of neuropeptides from specific sensory neurons ^86^. Our findings show that in addition to the transcriptional regulation of individual insulin genes during starvation conditions, a remodeling of the entire secretion machinery of the gut occurs during acute nutritional deprivation, indicating a broader role of the gut epithelia as a secretory organ that controls animal behavior. This expands on the previously described ‘FoxO-to-FoxO’ signaling mechanism by demonstrating that intestinal DAF-16/FoxO modifies the function of other organs not just by mediating insulin-mediated cross-tissue FoxO activation ^46^, but also via coordinating other interorgan signaling axes that do not directly involve insulin signaling. An internal state-induced global block of all gut-to-brain signaling during adverse environments would affect not only the release of neuropeptides, but also of yolk proteins, metabolites and other signaling molecules that are released from the gut in secretory vesicles to modulate several aspects of the animal’s physiology ^87^.

We found that the pacemaker neurons in the ENS of the *C. elegans* foregut are not receptive to the gut-to-brain insulin signaling axis, which agrees with the previously proposed autonomous nature of the MC neurons ^23,25^. Instead, the neuromodulatory NSM neurons receive insulin cues from the midgut epithelial cells to modify the output from the foregut ENS circuit depending on the nutritional status of the animal. This is supported by a previous study that found an altered response of the NSM neurons to food stimuli in animals that have experienced extended food deprivation ^88^. Altered 5-HT release from NSM neurons can modulate the rate of foregut contractions by acting on known serotonergic receptors that are expressed in both neurons and muscles of the foregut, as described previously ^74,76,89^. For the non-insulin peptidergic signal NLP-40 from the midgut that regulates defecation behavior by activating hindgut enteric neurons ^19^, our results demonstrate a physiological regulation of the release of this neuropeptide during nutrient scarcity. A reduced rate of expulsion from the hindgut of a starved animal would increase the residence time of food in its intestinal lumen, which will likely support increased nutrient absorption, a phenomenon that has been observed when *Drosophila* is exposed to low nutrient conditions ^8^.

We observe different feeding responses in starved adults and dauer diapause arrested larvae on encountering food. This difference is likely due to the activity of an additional stress-responsive transcription factor, DAF-12/VDR, during dauer remodeling ^29^, which cell autonomously shuts down pacemaker activity in the ENS of the foregut in dauers ^39^.

Reconstituting the gut-to-pharynx insulin signaling during the dauer diapause stage activates this silenced circuit likely via the release of 5-HT from the NSM neurons, which is supported by a previous study that induced foregut contractions in dauer stage animals by exposing them to exogenous 5-HT ^90^. While the inhibitory effect on foregut contractions is unambiguous in the starvation-induced dauer diapause stage, the effect of prolonged starvation on the feeding rate of adults is dependent on the conditions in which the behavior is recorded. Initial studies reported an increase in foregut contractions in starved animals, but this behavior was observed in the absence of food ^26,91^. A subsequent study found an increase in the rate of feeding immediately after starved adults were returned to food ^27^. This is consistent with strong calcium activity in the NSM neurons after a starved animal encounters food and this activity lasts for less than five minutes after food exposure ^88^. These findings indicate a heightened sensitivity of the animal to food-derived cues after an extended period of starvation. Instead, if a starved animal is allowed to habituate on food for 5 minutes, then its rate of feeding is consistently lower than that in the fed state, which we have validated in blinded experiments (Figure S1A). Two recent studies that quantified feeding behavior over longer time scales, one using automated pharyngeal pumping measurements and another that quantified feeding rate independent of foregut contractions, have not observed an increase in the rate of feeding after starved animals are returned to food ^24,92^. This emphasizes the importance of separating the immediate response to the introduction of a new stimulus from the persistent effect of a change in the internal state of the animal in studies that measure behavioral output from autonomous circuits.

The NSM neurons in the foregut enteric circuit can modulate behavioral outputs by altering overall brain state, which involves volumetric release of 5-HT to modify neural activity across multiple circuits via extra-synaptic communication ^76,93^. Axonal arborization patterns of serotonergic neurons in the mammalian brain suggest that 5-HT release from individual neuron types can potentially have widespread effects on brain activity ^94^. In the context of enteric circuits, the role of neuronally-released 5-HT in enhancing gut contractions is evolutionarily conserved in the *Drosophila* and mammalian gut ^95,96^, but how these regulatory mechanisms are influenced during different physiological states of the animal need further exploration. Altered 5-HT signaling has been linked to functional gastrointestinal disorders (FGID) in several clinical studies, but the internal or external factors that lead to 5-HT dysregulation in these patients remain poorly understood ^97^. Understanding how impairment of insulin signaling affects the function of serotonergic neurons in the ENS might shed new light on this unexplored aspect of gastrointestinal physiology.

## Supporting information

Key Resources Table

## RESOURCE AVAILABILITY

### Lead contact

Requests for further information and resources should be directed to and will be fulfilled by the lead contact, Oliver Hobert.

### Materials availability

All new *C. elegans* strains generated in this study (Key Resources Table) will be made available at the Caenorhabditis Genetics Center (CGC).

### Data and code availability

- RNA-sequencing data generated in this study have been deposited at NCBI Gene Expression Omnibus (GEO) at GEO: GSE285886 and are publicly available as of the date of publication.
- This paper did not generate any original code.
- Any additional information required to reanalyze the data reported in this paper is available from the lead contact upon request.

## Acknowledgments

We thank Qi Chen for *C. elegans* microinjections, Patrick Hu for sharing *trap-1* strains, Itai A. Toker and Eyal Ben-David for sharing their unpublished single-cell RNA-seq dataset, Wen Xi Cao for sharing the *pha-4prom2* sequence and members of the Hobert lab for comments on the manuscript. Some strains were provided by the CGC, which is funded by NIH Office of Research Infrastructure Programs (P40 OD010440). O.H. is an Investigator with the Howard Hughes Medical Institute, which funded this work.

## Author contributions

Conceptualization, S.S. and O.H.; Methodology, S.S.; investigation, S.S. and Z.W.; formal analysis, S.S. and Z.W.; writing – original draft, S.S. and O.H.; writing – review & editing, S.S. and O.H.; supervision, O.H.; funding acquisition, O.H.

## SUPPLEMENTAL FIGURES

**Figure S1.**
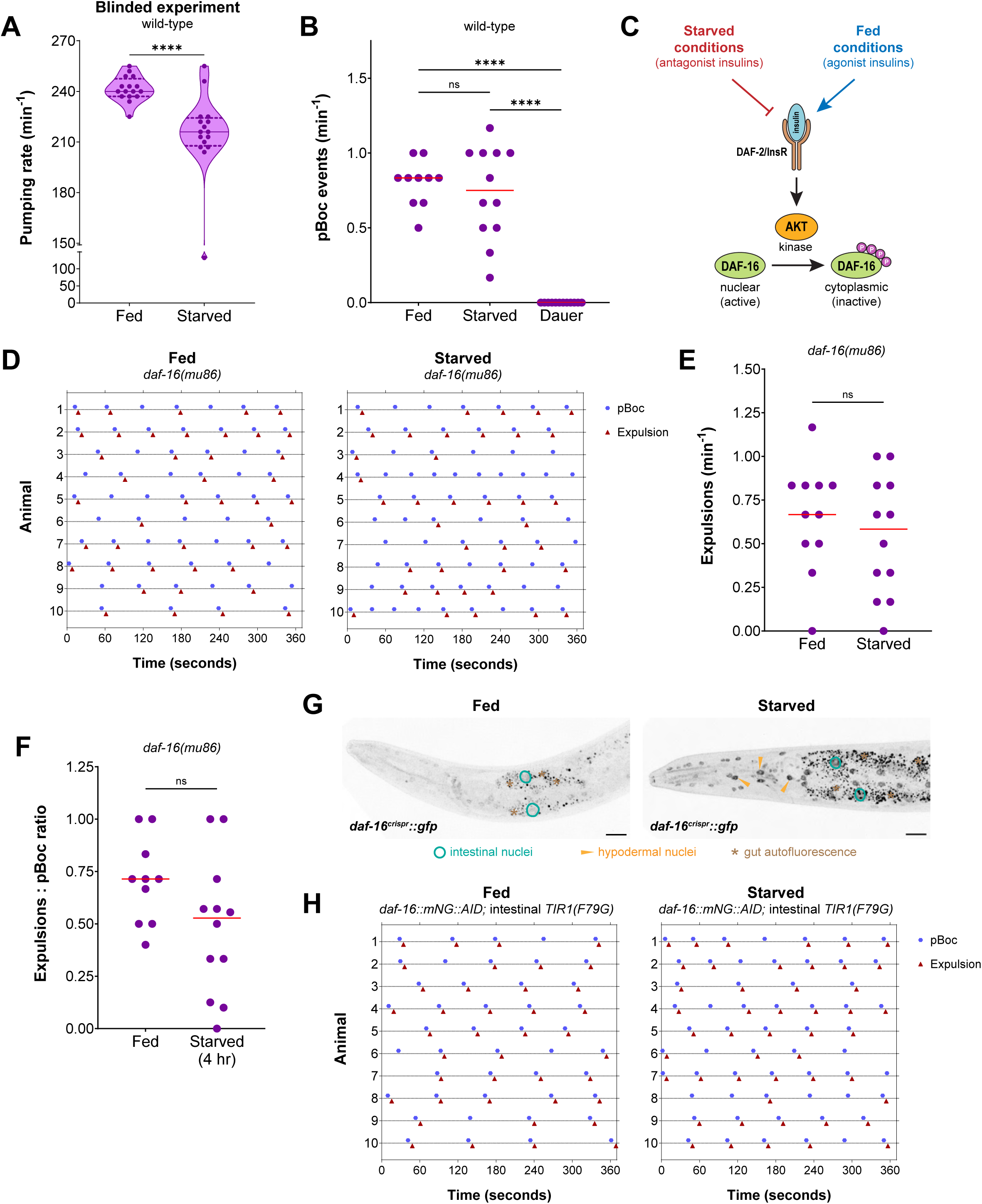
DAF-16/FoxO activity in the intestine non-cell autonomously regulates feeding and defecation behaviors, related to. Figure 1. (A) Pharyngeal pumping rate on food in fed vs starved (4 hr) wild-type adults in a blinded experiment. (B) Frequency of pBoc events on food in fed vs starved (4 hr) wild-type adults and in starvation-induced dauer stage animals. (C) Schematic of DAF-2/InsR signaling regulating DAF-16 nuclear localization in fed vs starved conditions. (D) Representative traces of pBoc and expulsion events on food in fed vs starved (4 hr) *daf-16(mu86)* adults (10 animals per condition). (E) Frequency of expulsion on food in fed vs starved (4 hr) *daf-16(mu86)* adults. (F) Expulsion: pBoc ratio on food in fed vs starved (4 hr) *daf-16(mu86)* adults. (G) Endogenous DAF-16 protein localization in fed vs starved (4 hr) *daf-16(ot971[daf-16::gfp])* adults. Representative images of 15 animals per condition. Scale bars, 20 μm. (H) Representative traces of pBoc and expulsion events on food after intestine-specific DAF-16 depletion in fed vs starved (4 hr) *daf-16(ot853); otSi2[ges-1p::TIR1(F79G)]* adults treated with 100 μM 5-Ph-IAA (10 animals per condition). Horizontal line in the middle of data points represents median value of biological replicates in (A, B, E, F). Additional horizontal lines represent 25^th^ and 75^th^ percentiles in (A). **** and ns represent *P* < 0.0001 and not significant, respectively, in Mann-Whitney test in (A, E, F) and in Dunn’s multiple comparison test after Kruskal-Wallis test in (B).

**Figure S2.**
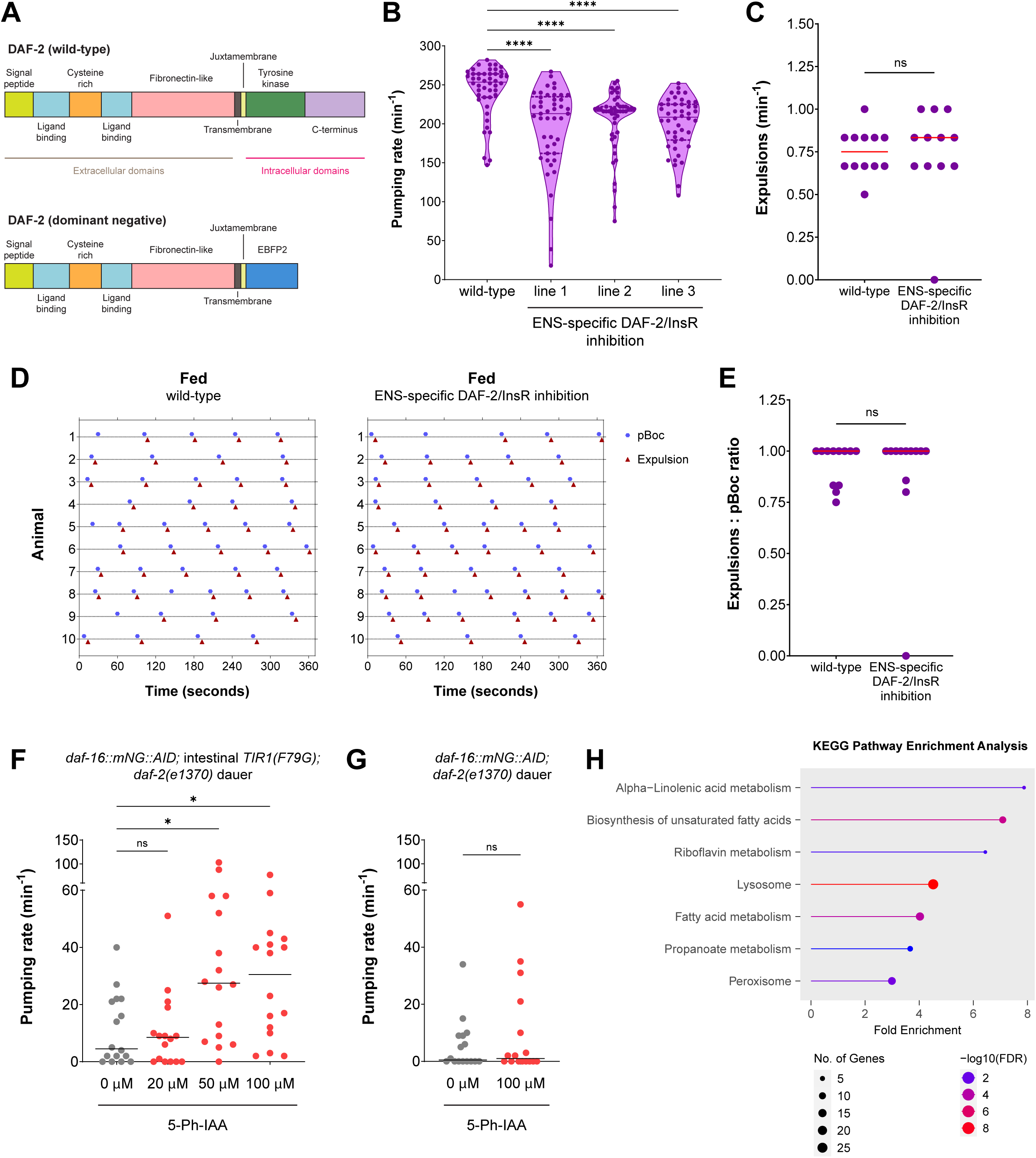
DAF-16/FoxO activity in the midgut regulates ENS output via modulating gut-to-ENS insulin signaling, related to. Figure 2. (A) Schematic showing the design of DAF-2(DN) construct in which the intracellular kinase domain of DAF-2/InsR is replaced by EBFP2 fluorescent protein. (B) Pharyngeal pumping rate on food in fed wild-type and *pha-4prom2::daf-2(DN)*-expressing (*otIs911*, *otIs912* and *otIs913* lines) adults. (C) Frequency of expulsion on food in fed wild-type and *otIs913[pha-4prom2::daf-2(DN)]* adults. (D) Representative traces of pBoc and expulsion (Exp) events on food in fed wild-type and *otIs913[pha-4prom2::daf-2(DN)]* adults (10 animals per condition). (E) Expulsion: pBoc ratio on food in fed wild-type and *otIs913[pha-4prom2::daf-2(DN)]* adults. (F) Pharyngeal pumping rate on food in *daf-16(ot853); otSi2[ges-1p::TIR1(F79G)]; daf-2(e1370)* dauer stage animals treated with either solvent (ethanol) or different concentrations of 5-Ph-IAA. (G) Pharyngeal pumping rate on food in *daf-16(ot853); daf-2(e1370)* dauer stage animals treated with either solvent (ethanol) or 100 μM 5-Ph-IAA. (H) KEGG pathway enrichment analysis of genes that are upregulated after intestine-specific DAF-16 depletion in dauer stage animals. Horizontal line in the middle of data points represents median value of biological replicates in (B, C, E-G). Additional horizontal lines represent 25^th^ and 75^th^ percentiles in (B). *, **** and ns represent *P* < 0.05, *P* < 0.0001 and not significant, respectively, in Dunn’s multiple comparison test after Kruskal-Wallis test in (B, F) and in Mann-Whitney test in (C, E, G).

**Figure S3.**
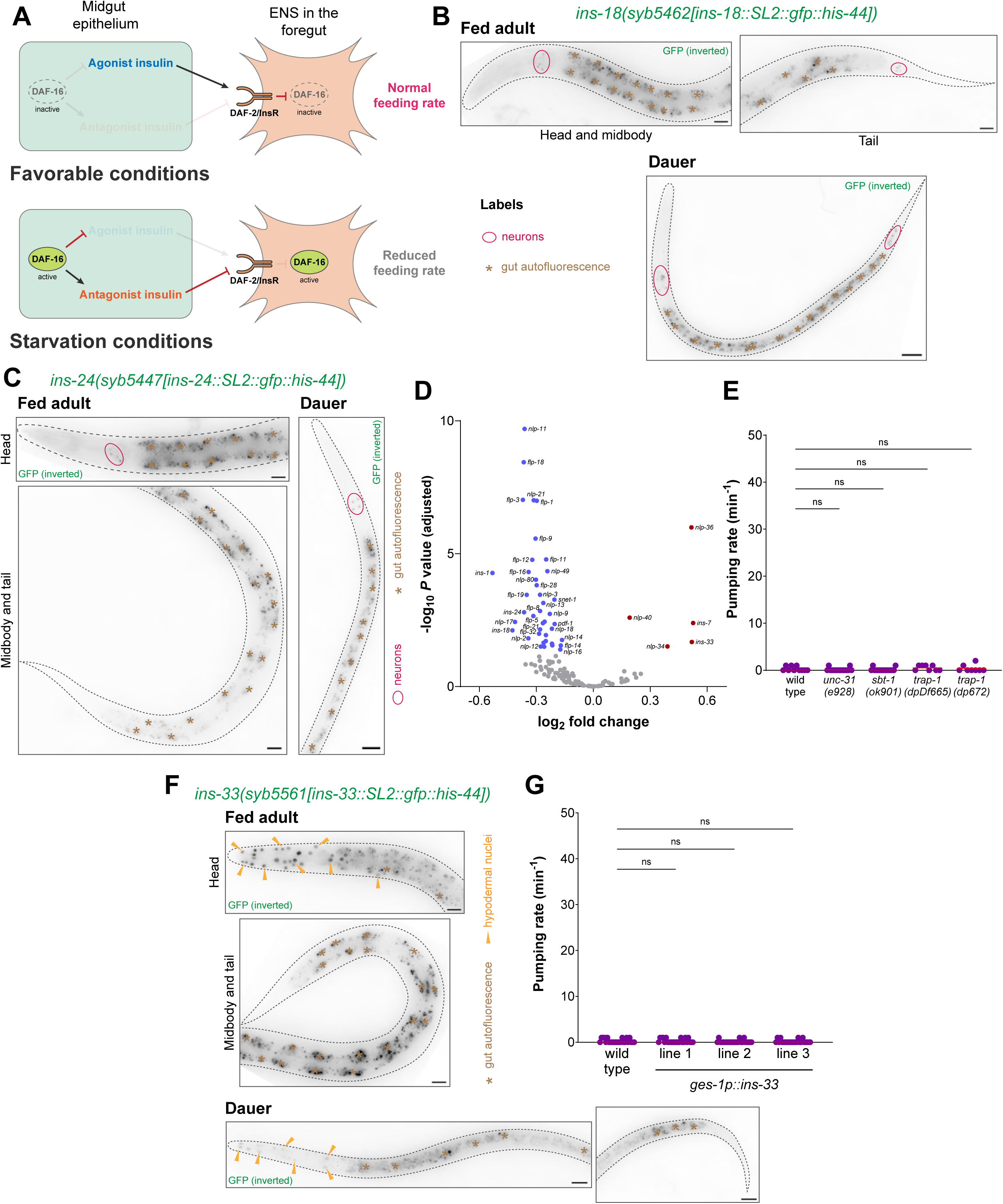
DAF-16/FoxO activity in the dauer intestine changes the expression of insulin genes both autonomously and non-cell autonomously, related to. Figure 3. (A) Two alternative models for insulin peptides released from the intestine regulating output from the ENS in the foregut – (1) absence of agonist insulin or (2) release of antagonist insulin reduces the rate of foregut contractions in starvation conditions. (B) Expression of the endogenously tagged *ins-18* reporter allele *syb5462[ins-18::SL2::gfp::his-44]* in fed adult and starvation-induced dauer stage animals. Representative images of 15 animals per condition. Scale bars, 20 μm. (C) Expression of the endogenously tagged *ins-24* reporter allele *syb5447[ins-24::SL2::gfp::his-44]* in fed adult and starvation-induced dauer stage animals. Representative images of 15 animals per condition. Scale bars, 20 μm. (D) Volcano plot for all neuropeptide family genes that are differentially expressed after intestine-specific DAF-16 depletion in dauers. (E) Pharyngeal pumping rate on food in wild type, *unc-31(e928)*, *sbt-1(ok901)*, *trap-1(dpDf665)* and *trap-1(dp672)* starvation-induced dauer stage animals. (F) Expression of the endogenously tagged *ins-33* reporter allele *syb5561[ins-33::SL2::gfp::his-44]* in fed adult and starvation-induced dauer stage animals. Representative images of 15 animals per condition. Scale bars, 20 μm. (G) Pharyngeal pumping rate on food in starvation-induced dauer stage animals that constitutively express *ins-33* in the intestine (*otEx8326*, *otEx8327* and *otEx8328*). Horizontal line in the middle of data points represents median value of biological replicates in (E, G). ns represents not significant in Dunn’s multiple comparison test after Kruskal-Wallis test in (E, G).

**Figure S4.**
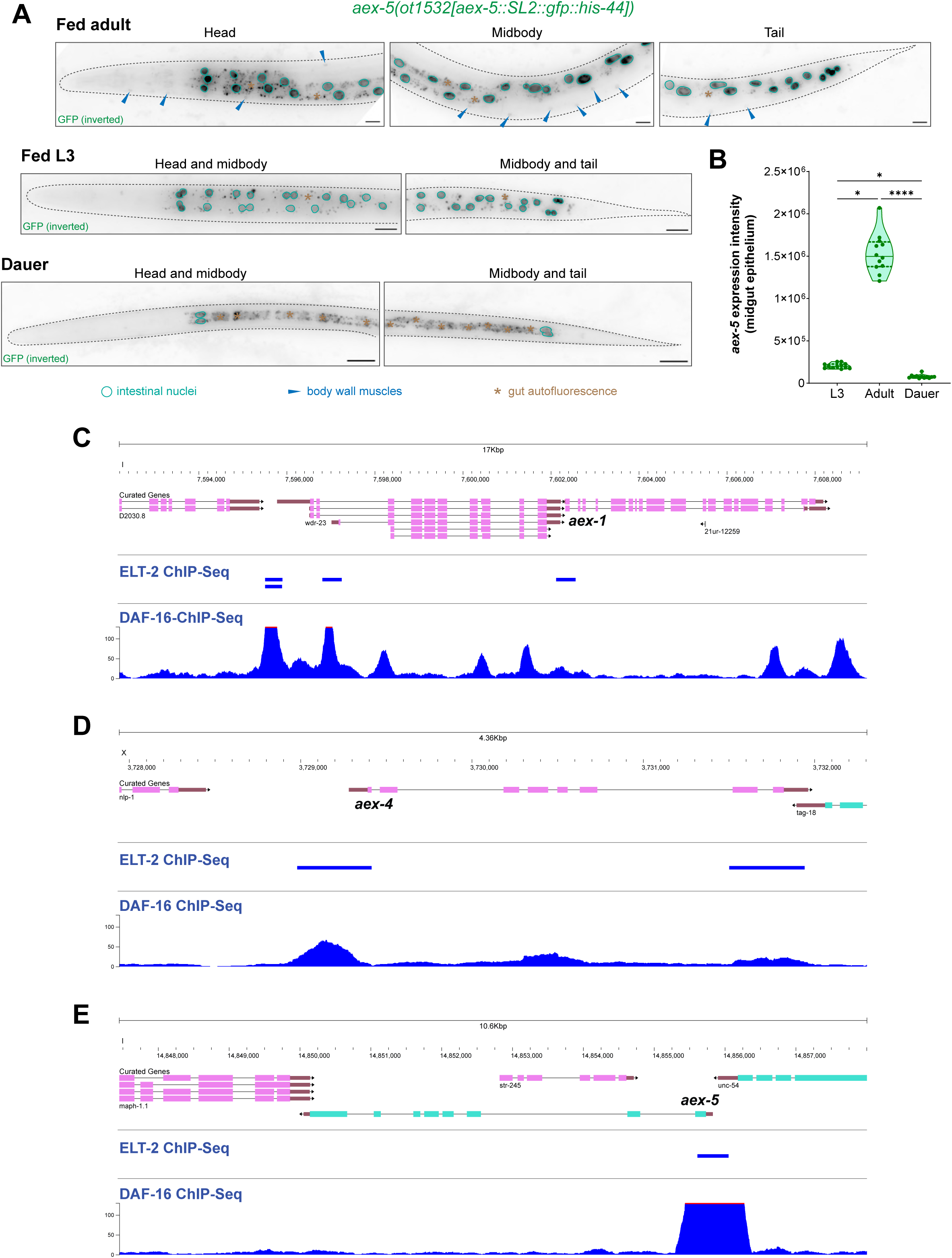
Multiple components of the neuropeptide secretion machinery of the gut are downregulated during acute starvation, related to. Figure 4. (A) Expression of the endogenously tagged *aex-5* reporter allele *aex-5(ot1532[aex-5::SL2::gfp::his-44])* in fed adult and L3 larva, and in starvation-induced dauer stage animals. Scale bars, 20 μm. (B) Quantification of *aex-5* expression in midgut epithelial cells of fed adult and L3 larva, and starvation-induced dauer stage animals. Horizontal line in the middle of data points and additional horizontal lines represent median of biological replicates, and 25^th^ and 75^th^ percentiles, respectively. * and **** and ns represent *P* < 0.05, *P* < 0.0001 and not significant, respectively, in Dunn’s multiple comparison test after Kruskal-Wallis test. (C) Location of ChIP-seq peaks for ELT-2 and DAF-16 in the genomic locus of *aex-1*. (D) Location of ChIP-seq peaks for ELT-2 and DAF-16 in the genomic locus of *aex-4*. (E) Location of ChIP-seq peaks for ELT-2 and DAF-16 in the genomic locus of *aex-5*.

**Figure S5.**
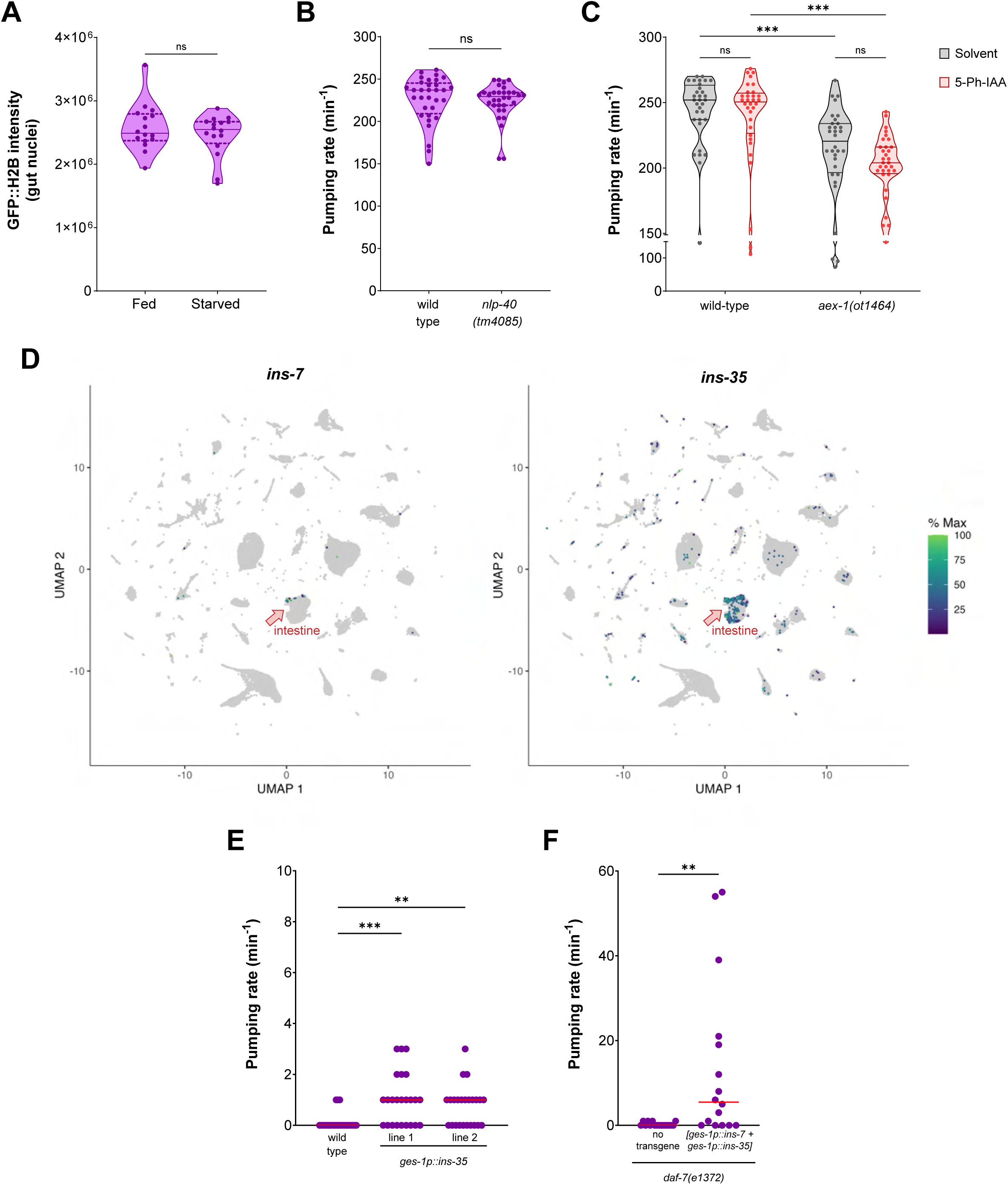
Secretion of gut-derived insulins is both necessary and sufficient for foregut contractility, related to. Figures 5 and **6**. (A) Quantification of GFP::H2B intensity in midgut nuclei of fed vs starved (24 hr) adults expressing *otIs904[ges-1p::ins-1::tagRFP::SL2::gfp::his-44]*. (B) Pharyngeal pumping rate on food in fed wild-type and *nlp-40(tm4085)* adults. (C) Pharyngeal pumping rate on food in fed *daf-16(ot853)* adults without *TIR1(F79G)* in wild-type or *aex-1(ot1464)* genetic background. Animals were treated with either solvent (ethanol) or 100 μM 5-Ph-IAA. (D) Intestine-specific expression of *ins-7* and *ins-35* retrieved from a single-cell RNA-seq dataset ^70^. (E) Pharyngeal pumping rate on food in starvation-induced dauer stage animals that constitutively express *ins-35* in the intestine (*otEx8242* and *otEx8243*). (F) Pharyngeal pumping rate on food in *daf-7(e1372)* dauer stage animals that constitutively co-express *ins-7* and *ins-35* in the intestine (*otEx8223*). Horizontal line in the middle of data points represents median value of biological replicates in (A-C, E, F). Additional horizontal lines represent 25^th^ and 75^th^ percentiles in (A-C). **, *** and ns represent *P* < 0.01, *P* < 0.001 and not significant, respectively, in Mann-Whitney test in (A, B, F), in Sidak’s multiple comparisons test after two-way ANOVA in (C) and in Dunn’s multiple comparison test after Kruskal-Wallis test in (E).

### Supplemental information

**Table S1:** Differentially expressed genes after intestinal DAF-16 depletion in dauers, related to Figure 2. List of all genes that are differentially expressed after intestine-specific DAF-16 removal in dauers using AID2, in comparison to either no TIR1 or no 5-Ph-IAA control.

**Table S2:** Enrichment of transcription factor binding sites in promoter regions of genes differentially expressed after intestinal DAF-16 depletion in dauers, related to Figure 2.

**Table S3:** crRNA and single-stranded oligodeoxynucleotide sequences used for CRISPR/Cas9-mediated genome editing, related to STAR Methods.

## STAR METHODS

### EXPERIMENTAL MODEL AND STUDY PARTICIPANT DETAILS

#### Maintenance of *C. elegans* strains

All *C. elegans* strains were maintained at 20°C on nematode growth medium (NGM) plates seeded with *Escherichia coli* (OP50 strain), unless otherwise specified. Strains with temperature-sensitive *daf-2(e1370)* and *daf-7(e1372)* alleles were maintained at 15°C. All strains were grown in optimum conditions in the abundance of food for at least three generations prior to using them in any experiment. All experiments in this study were performed using animals that were grown at 25°C for their entire developmental period (from unhatched embryos to adulthood) to be consistent with the temperature used for starvation-induced dauer formation assays.

## METHOD DETAILS

### DNA constructs and microinjections

All plasmids used in this study were generated via NEBuilder HiFi DNA assembly using the manufacturer’s protocol. All plasmid sequences were validated using Sanger (Azenta) and/or Oxford Nanopore (Plasmidsaurus) sequencing. The *pha-4prom2* sequence in pSS35 [*pha-4prom2::TIR1(F79G)::mTur2::tbb-2 3’ UTR*] and pSS36 [*pha-4prom2::daf-2(DN)::ebfp2::tbb-2 3’ UTR*] is a 2,393 bp fragment upstream of isoform *b* of the *pha-4* gene (−2,387 to +6 position with respect to the start codon of *pha-4* isoform *b*), which drives expression in all 22 enteric neurons (20 pharyngeal neurons and the two hindgut-innervating neurons, AVL and DVB) and also in RIS and PVT neurons. The *TIR1(F79G)* sequence in pSS35 was generated by introducing the *F79G* [Phe(TTC) to Gly(GGA)] mutation in the *Arabidopsis thaliana TIR1* sequence, as described previously ^44^. The *daf-2(DN)::ebfp2* sequence in pSS36 is from a previous study that replaced the tyrosine kinase and carboxyl-terminal domains of the cDNA of *daf-2* isoform *a* (corresponding to amino acids 1,246 to 1,846) with the blue fluorescent protein *ebfp2* sequence^39^. The *tbb-2 3’ UTR* sequence is a 156 bp sequence immediately downstream of the stop codon of the *tbb-2* gene. The *ceh-19prom2* sequence of pSS43 [*ceh-19prom2::daf-2(DN)::eBFP2::SL2::tagRFP-T::tbb-2 3’ UTR*] is a 1,522 bp fragment upstream of isoform *b* of the *ceh-19* gene (−1,516 to +6 position with respect to the start codon of *ceh-19* isoform *b*). The *tph-1prom5* sequence of pSS44 [*tph-1prom5::daf-2(DN)::eBFP2::SL2::tagRFP-T::tbb-2 3’ UTR*] is a previously described 232 bp fragment upstream of isoform *a* of the *tph-1* gene (−232 to −1 position with respect to the start codon of *tph-1* isoform *a*) ^75^.

The pSS29 [*ges-1p::ins-1::tagRFP-T::SL2::gfp::his-44::tbb-2 3’ UTR*] and pSS34 [*ges-1p::ins-1::tagRFP-T::SL2::ebfp2::his-44::tbb-2 3’ UTR*] constructs were generated by cloning the entire genomic locus (start codon to stop codon) of the *ins-1* gene downstream of a 1,999 bp *ges-1* promoter fragment (−1,999 to −1 position with respect to the start codon of *ges-1*). The pSS31 [*ges-1p::nlp-40::tagRFP-T::SL2::gfp::his-44::tbb-2 3’ UTR*] construct was generated by cloning the cDNA of isoform *a* of the *nlp-40* gene downstream of the *ges-1* promoter fragment described above. pZW1 [*ges-1p::ins-7::tbb-2 3’ UTR*], pZW3 [*ges-1p::ins-33::tbb-2 3’ UTR*] and pZW24 [*ges-1p::ins-35::tbb-2 3’ UTR*] were generated by cloning the entire genomic locus (start codon to stop codon) of *ins-7* isoform *b*, *ins-33* and *ins-35*, respectively, downstream of the *ges-1* promoter fragment described above.

Before microinjection, all plasmids were linearized using a single cutter restriction enzyme that cuts in the plasmid backbone. Linearized plasmids were injected at the concentrations listed in the Key Resources Table into both gonadal arms of young adult animals. F1 progeny was picked based on expression of the co-injection marker and animals that transmitted the array to F2 progeny were used to generate transgenic lines. At least two independent lines (obtained from different injected P0 animals) for each injection type were used in subsequent experiments.

### CRISPR/Cas9 genome editing

CRISPR/Cas9 genome editing was performed using a modified version of a previously described protocol ^99^. All CRISPR injection mixes constituted 250 ng/μL of *S. pyogenes* Cas9 nuclease (IDT), 100 ng/μL of tracrRNA (IDT), 56 ng/μL (combined) of all crRNAs (IDT) and 100 ng/μL of each single-stranded oligodeoxynucleotide (ssODN) repair template (IDT). First, tracrRNA and all crRNAs were incubated at 95°C for 5 min followed by at 10°C for 5 min. Cas9 was added to this mix, pipetted several times and incubated at 25°C for 10 min. All ssODN and plasmids were subsequently added to this mix and the volume was filled to 20 μL using nuclease-free water. The mixture was centrifuged at 17,900 xg for 2 min and the supernatant was used for microinjections. For all CRISPR edits on chromosome II, the pRF4 [*rol-6p::rol-6(su1006)*] plasmid was co-injected at a concentration of 60 ng/μL. For CRISPR edits on all other chromosomes, *dpy-10* co-CRISPR was performed using crRNA and ssODN listed in Table S3. For all CRISPR edits, F1 animals were picked on the basis of roller phenotype and were genotyped by PCR amplification of the edited locus followed by Sanger (Azenta) or Oxford Nanopore (Plasmidsaurus) sequencing. For confirmed heterozygous edits, non-roller progeny was singled from F1 plates and genotyped by PCR to obtain homozygous lines.

The genomic sequences for all genes edited in this study were obtained from WormBase^100^. The *otSi2[ges-1p::TIR1(F79G)::mRuby::unc-54 3’ UTR]* single-copy transgene was generated using a previously described technique by introducing the *F79G* [Phe(TTC) to Gly(GGA)] mutation in the *TIR1* sequence of *ieSi61[ges-1p::TIR1::mRuby::unc-54 3’ UTR]* using a crRNA and an ssODN listed in Table S3 ^44^. The *ins-18(ot1326)*, *ins-18(ot1328)*, *ins-1(ot1360)*, *ins-1(ot1363)*, *aex-1(ot1357)*, *aex-1(ot1464)*, *ins-7(ot1427)*, *ins-35(ot1443)* and *tph-1(ot1545)* alleles were obtained by deleting the entire coding sequence (start codon to stop codon) of the corresponding gene using two crRNAs and an ssODN listed in Table S3. The *otDf2* allele was generated using two crRNAs and an ssODN listed in Table S3 to delete a 25.5 kb cluster of insulin genes, many of which (*ins-24*, *ins-28*, *ins-29* and *ins-30*) are upregulated in dauers ^48^.

The *aex-1(ot1543[aex-1::SL2::gfp::his-44])*, *aex-4(ot1530[aex-4::SL2::gfp::his-44])* and *aex-5(ot1532[aex-5::SL2::gfp::his-44])* alleles were generated using a crRNA (listed in Table S3) and a long single-stranded repair template to insert the *SL2::gfp::his-44* sequence immediately after the stop codon of the corresponding gene. For *aex-1(ot1543[aex-1::SL2::gfp::his-44])*, the *SL2::gfp::his-44* sequence was inserted after the stop codon of isoform *a* of *aex-1*. The long single-stranded repair templates were generated using a previously described method ^99^. The *SL2::gfp::his-44* sequence was PCR amplified with primers containing 35 bp homology sequence corresponding to each end of the genomic locus to be edited. One of the two primers used for PCR amplification had a 5’ phosphate modification. 10 μg of the PCR product was digested with lambda exonuclease (NEB) to digest the DNA strand with 5’ phosphorylation. The unphosphorylated strand of DNA was purified using Monarch PCR & DNA Cleanup Kit (NEB) and was used as a repair template at a final concentration of 100 ng/μL in the CRISPR injection mix.

Suny Biotech generated the *ins-1(syb5452[ins-1::SL2::gfp::his-44])*, *ins-18(syb5462[ins-18::SL2::gfp::his-44])*, *ins-24(syb5447[ins-24::SL2::gfp::his-44])*, *ins-7(syb5424[ins-7::SL2::gfp::his-44])*, *ins-33(syb5561[ins-33::SL2::gfp::his-44])* and *ins-35(syb6236[ins-35::SL2::gfp::his-44])* alleles by inserting the *SL2::gfp::his-44* sequence immediately after the stop codon of the corresponding gene.

### FLInt-mediated transgene integration

The *otIs904*, *otIs908*, *otIs911*, *otIs912*, *otIs913*, *otIs927*, *otIs937* and *otIs942* transgenes were inserted at the intergenic *oxTi553[eft-3p::tdTomato::H2B::unc-54 3’ UTR + Cbr-unc-119(+)] V.* locus using the Fluorescent Landmark Interference (FLInt) method ^101^. The FLInt injection mixes were prepared similar to CRISPR mixes, i.e., first, tracrRNA (final concentration of 100 ng/μL) and a crRNA (final concentration: 56 ng/μL) targeting the tdTomato locus (listed in Table S3) were incubated at 95°C for 5 min followed by at 10°C for 5 min. Cas9 (final concentration: 250 ng/μL) was added to this mix, pipetted several times and incubated at 25°C for 10 min. After this, all plasmids were added to the mix at the concentrations listed in the Key Resources Table. The volume was filled to 20 μL using nuclease-free water, centrifuged at 17,900 xg for 2 min and the supernatant was used for microinjections. All injected animals were maintained at room temperature and F1 progeny expressing the co-injection marker were singled onto plates. F2 progeny was singled from F1 plates that had at least 75% of the animals expressing the co-injection marker. F2 plates that transmitted the co-injection marker to 100% of the progeny were used to generate lines.

### 5-Ph-IAA treatment for AID2

For all AID2 experiments except the RNA-seq protocol, the animals were treated with the desired concentration of 5-phenyl-indole-3-acetic acid (5-Ph-IAA) as described previously with some modifications ^102^. A 100 μL freshly prepared solution of 5-Ph-IAA in 25% ethanol (v/v) was added on top of 60 mm NGM plates seeded with OP50 bacteria. For control plates, an equal volume of 25% ethanol (v/v) was added such that the final ethanol concentration on the plate is 0.25% (v/v). All plates were stored in the dark for the entire duration of the experiment. Age-synchronized embryos obtained via the alkaline bleaching method were transferred to the plates one day after addition of 5-Ph-IAA and the animals were grown at 25°C till they reached the desired developmental stage for the experiment.

### Dauer formation assays

A previously described protocol was used to obtain starvation-induced dauers ^103^. Eight gravid adult worms were transferred to NGM plates seeded with OP50 bacteria at 25°C. After 6-8 days when all animals on the plates were starved due to lack of bacterial food, animals were collected in 1% SDS and incubated for 20 minutes with gentle agitation to kill all non-dauer stages. Animals were centrifuges at 1,150 xg for 2 min and washed twice with M9 buffer to remove all SDS. At this point, the tube contained living dauer stage animals and the carcasses of non-dauer stages. Animals were pipetted on a fresh NGM plate seeded with OP50 bacteria and live dauer animals were used for experiments within 40 min of removing the 1% SDS solution.

Dauers for strains with the *daf-2(e1370)* or *daf-7(e1372)* mutation were generated by transferring age-synchronized embryos obtained via the alkaline bleaching method to NGM plates seeded with OP50 bacteria. Animals were maintained at 25°C for three days, after which all animals carrying the *daf-2(e1370)* or *daf-7(e1372)* mutation were arrested in the dauer stage.

### Adult starvation assays

To starve adult animals, age-synchronized embryos obtained via the alkaline bleaching method were transferred to NGM plates seeded with OP50 bacteria at 25°C. After 48 hours, adult stage animals were washed off the plate using M9 buffer and centrifuged at 300 xg for 1 min. Animals were then washed three times with M9 buffer to get rid of all bacteria. For the first two washes, the supernatant was removed after centrifugation at 300 xg for 1 min. For the final wash, animals were allowed to settle down by gravity for 5 min after which the supernatant was removed, and animals were pipetted on unseeded NGM plates for the desired amount of time. At the end of the starvation period at 25°C, animals were transferred to NGM plates seeded with OP50 bacteria and were used for experiments within 15 min of transferring them on food.

### Pharyngeal pumping assays

All pharyngeal pumping assays were performed on freely moving animals. The required number of animals were transferred to an NGM plate seeded with a uniform thin layer of OP50 bacteria. Animals were allowed to settle down for 5 min and the movement of the grinder of the pharynx was recorded using a hand-held tally counter by observing the animals under a Nikon Eclipse E400 upright microscope equipped with DIC optics. For age-synchronized adults, the number of grinder movements in a 20 sec period was recorded using a 20x air objective lens of the Nikon Eclipse E400 microscope and was multiplied by three to obtain pharyngeal pumps per minute. For dauer stage animals, the number of grinder movements in a 1 min period was recorded using a 50x air objective lens of the Nikon Eclipse E400 microscope. The rate of pharyngeal pumping was recorded from at least 8 animals on each day and at least on two independent days.

### Defecation assays

All defecation assays were performed on freely moving animals. The required number of animals were transferred to an NGM plate seeded with a uniform thin layer of OP50 bacteria. Animals were allowed to settle down for 5 min, after which the posterior end of animals is observed using a 20x air objective lens of a Nikon Eclipse E400 upright microscope equipped with DIC optics. Each animal was observed for a 6 min period, during which the timings of contraction of posterior body muscles (pBoc) and expulsion (Exp) muscle contraction that releases food contents from the anal opening were recorded using the Behavioral Observation Research Interactive Software (BORIS) software version 8.23 ^104^. Defecation behavior was recorded from at least 5 animals on each day and at least on two independent days.

### RNA isolation and library preparation

OH14654: *daf-16(ot853[daf-16::mNG::AID]); daf-2(e1370)* and OH17582: *daf-16(ot853[daf-16::mNG::AID]); otSi2[ges-1p::TIR1(F79G)::mRuby::unc-54 3’ UTR]; daf-2(e1370)* strains were used for this experiment. Prior to growing animals, 250 μL of a freshly prepared solution of 5-Ph-IAA in 25% ethanol (v/v) was uniformly spread on top of each unseeded 100 mm NGM plate to obtain a final 5-Ph-IAA concentration of 100 μM on the plates. For control plates, an equal volume of 25% ethanol (v/v) was added such that the final ethanol concentration on the plate is 0.2% (v/v). All plates were stored in the dark for the entire duration of the experiment. After one day of adding 5-Ph-IAA to the plates, a uniform thin layer of OP50 bacteria was spread on top of each plate and incubated at 25°C for an additional day. Subsequently, ∼8,000 age-synchronized embryos obtained via the alkaline bleaching method were transferred to each plate and on three plates per condition. All plates were maintained at 25°C for three days after which all animals were arrested in the dauer stage due to presence of the *daf-2(e1370)* mutation. Any animals that were not arrested in the dauer stage (less than 10 animals per plate) were picked and removed using a worm pick. A total of ∼24,000 animals from three 100 mm NGM plates for each condition, which constituted a biological replicate, were collected in M9 buffer. Animals were washed three times using M9 buffer with centrifugation at 1,300 xg for 2 min after each wash to remove all bacteria. 200 μL of TRIzol reagent (Thermo Fisher) was added to the worm pellet and samples were homogenized using a Kontes Pellet Pestle Motor (Fisher Scientific) for 30 seconds. Samples were immediately frozen on dry ice for 5 min and the homogenization-freezing cycles were repeated for a total of five times. All homogenized samples were stored at −80°C.

After five biological replicates were collected for each condition, total RNA was isolated from all samples using the RNeasy Micro Kit (Qiagen) using the manufacturer’s protocol. 100 ng of total RNA per sample was used for library preparation using the Universal RNA-seq with NuQuant kit (Tecan Genomics) using the manufacturer’s protocol with custom AnyDeplete (IC0149S) to deplete rRNA. All 20 libraries (4 conditions x 5 biological replicates) were pooled at a final concentration of 2 nM and were multiplexed for 75 bp single-end sequencing on a NextSeq 550 system (Illumina).

### RNA-seq data analysis and visualization

All sequencing analyses were performed on the Galaxy web platform ^105^. Quality control of raw RNA-seq reads were performed using FastQC. Adapter sequences were trimmed using Trimmomatic ^106^. The trimmed reads were aligned to the *C. elegans* genome (WS284 release) using STAR ^107^. Read counts were assigned to genes using featureCounts ^108^. Differential expression of genes across the four conditions was determined using DESeq2 ^109^. The principal component analysis (PCA) and sample-to-sample distance matrix plots were generated using DESeq2. The heatmap for differentially expressed genes was generated on the Galaxy web platform using the heatmap.2 function from the ggplot2 package on R.

The overlap between different gene lists were identified using Venny 2.1.0. Proportioned Venn diagrams were generated using Venn Diagram Plotter (Pacific Northwest National Laboratory). KEGG pathway enrichment analysis and enrichment of transcription factor binding motifs in upstream sequences of differentially expressed genes were performed using ShinyGO^110–112^. ChIP-seq peaks for DAF-16 and ELT-2 were visualized using JBrowse2 on Wormbase ^100^.

### Microscopy

Animals were mounted on a 5% agarose pad on top of a glass slide and were anesthetized using 50 mM sodium azide. Strains expressing the *daf-16(ot971[daf-16::gfp])* allele were anesthetized using 4 mM levamisole (instead of sodium azide). Animals expressing *daf-16(ot853)*, *daf-16(ot971)*, *otEx8059*, *otIs937* and *otIs942* were imaged on an inverted Zeiss LSM 980 laser scanning confocal microscope using a 40x water immersion objective lens. All other strains were imaged on a Zeiss Imager Z2 upright microscope equipped with Colibri 7 LEDs (Zeiss) using a 40x oil immersion objective lens. A full z-stack was acquired for all animals to cover their entire body width.

## QUANTIFICATION AND STATISTICAL ANALYSIS

### Image quantification

All images were processed and quantified on the Fiji platform on ImageJ ^113^. The mean background subtracted signal intensity from the two coelomocytes in the anterior region of the body was quantified for INS-1::TagRFP and NLP-40::TagRFP secretion assays. For the same assays, the signal intensity from the intestinal lumen and intestinal cytoplasm were calculated as the mean background subtracted intensity from two different regions of the intestinal lumen or cytoplasm, respectively. For quantification of signal intensity from the intestinal nuclei, the mean background subtracted intensity was calculated from two intestinal nuclei, one each from the first two rows of the intestinal cells.

### Statistical analysis

Data points for all biological replicates are displayed in the figures. Statistical tests used in each figure and the corresponding *P* values are listed in the figure legends. *P* < 0.05 was considered to be statistically significant. To calculate the *P* value for overlap between two gene lists, hypergeometric tests were performed on R. All other statistical tests and plotting were performed on GraphPad Prism 10.

For each type of quantitative assay described in this study, sample sizes were estimated by first performing a pilot experiment using eight animals per condition to determine the mean and standard deviation. Using these values, the required sample size was estimated using G*Power for a type I error rate of 0.05 and a statistical power of 0.95 ^114^. If the estimated sample size per condition was less than 35, the experiment was performed again using the estimated total number of animals. For all subsequent assays of the same type, the same sample size per condition was used.

