## Supplementary material for "Gut epithelium modifies enteric behaviors during nutritional adversity via distinct peptidergic signaling axes": Key Resources Table

| REAGENT or RESOURCE | SOURCE | IDENTIFIER |
|---|---|---|
| Bacterial and virus strains | | |
| *Escherichia coli* OP50 strain | Caenorhabditis Genetics Center (CGC) | https://cgc.umn.edu/strain/OP50 |
| Chemicals, peptides, and recombinant proteins | | |
| Alt-R *S. pyogenes* Cas9 nuclease V3 | Integrated DNA Technologies | Catalog No. 1081059 |
| Lambda exonuclease | New England Biolabs | Catalog No. M0262S |
| 5-phenyl-indole-3-acetic acid (5-Ph-IAA) | BioAcademia | Catalog No. 30-003-10 |
| Critical commercial assays | | |
| NEBuilder HiFi DNA Assembly Master Mix | New England Biolabs | Catalog No. E2621L |
| Monarch PCR & DNA Cleanup Kit | New England Biolabs | Catalog No. T1030 |
| TRIzol reagent | Thermo Scientific | Catalog No. 15596026 |
| RNeasy Micro Kit | Qiagen | Catalog No. 74004 |
| Universal RNA-seq with NuQuant kit, AnyDeplete IC0149S | Tecan Genomics | Catalog No. 0364 |
| Deposited data | | |
| Whole-dauer RNA-seq data | This paper | GEO: GSE285886 |
| Experimental models: Organisms/strains | | |
| *Caenorhabditis elegans* wild-type N2 strain | Caenorhabditis Genetics Center (CGC) | https://cgc.umn.edu/strain/N2 |
| *daf-16(mu86) I.* | Caenorhabditis Genetics Center (CGC) | https://cgc.umn.edu/strain/CF1038 |
| *daf-16(ot853[daf-16::mNG::AID]) I; otSi2[ges-1p::TIR1(F79G)::mRuby::unc-54 3' UTR *ieSi61] II; daf-2(e1370) III.* | This paper | OH17582 |
| *daf-16(ot853[daf-16::mNG::AID]) I; otSi2[ges-1p::TIR1(F79G)::mRuby::unc-54 3' UTR *ieSi61] II.* | This paper | OH19232 |
| *daf-16(ot971[daf-16::gfp]) I.* | PMID: 32550509 | OH16024 |
| *otIs911[pha-4prom2::daf-2(DN)::ebfp2::tbb-2 3' UTR (10 ng/μL), unc-122p::mCherry::unc-54 3' UTR (5 ng/μL), GeneRuler 1 kb plus DNA ladder (100 ng/μL)] V.* | This paper | OH19081 |
| *otIs912[pha-4prom2::daf-2(DN)::ebfp2::tbb-2 3' UTR (10 ng/μL), unc-122p::mCherry::unc-54 3' UTR (5 ng/μL), GeneRuler 1 kb plus DNA ladder (100 ng/μL)] V.* | This paper | OH19082 |
| *otIs913[pha-4prom2::daf-2(DN)::ebfp2::tbb-2 3' UTR (10 ng/μL), unc-122p::mCherry::unc-54 3' UTR (5 ng/μL), GeneRuler 1 kb plus DNA ladder (100 ng/μL)] V.* | This paper | OH19118 |
| *daf-16(ot853[daf-16::mNG::AID]) I; daf-2(e1370) III.* | PMID: 33891586 | OH14654 |
| *ins-1(syb5452[ins-1::SL2::gfp::his-44]) IV.* | This paper | PHX5452 |
| *ins-18(syb5462[ins-18::SL2::gfp::his-44]) I.* | This paper | PHX5462 |
| *ins-24(syb5447[ins-24::SL2::gfp::his-44]) I.* | This paper | PHX5447 |
| *daf-16(ot971[daf-16::gfp]) ins-24(otDf2) I.* | This paper | OH17064 |
| *ins-18(ot1326) daf-16(ot971[daf-16::gfp]) I.* | This paper | OH18320 |
| *ins-18(ot1328) daf-16(ot971[daf-16::gfp]) ins-24(otDf2) I.* | This paper | OH18322 |

| | | |
|---|---|---|
| *daf-16(ot971[daf-16::gfp]) I; ins-1(ot1360) IV.* | This paper | OH18508 |
| *ins-18(ot1328) daf-16(ot971[daf-16::gfp]) ins-24(otDf2) I; ins-1(ot1363) IV.* | This paper | OH18511 |
| *unc-31(e928) IV.* | Caenorhabditis Genetics Center (CGC) | https://cgc.umn.edu/strain/DA509 |
| *sbt-1(ok901) V.* | Caenorhabditis Genetics Center (CGC) | https://cgc.umn.edu/strain/RB987 |
| *trap-1(dpDf665) I.* | Patrick Hu, PMID: 31840061 | OH16568 |
| *trap-1(dp672) I.* | Patrick Hu, PMID: 31840061 | OH16567 |
| *aex-1(sa9) I.* | Caenorhabditis Genetics Center (CGC) | https://cgc.umn.edu/strain/JT9 |
| *aex-5(sa23) I.* | Caenorhabditis Genetics Center (CGC) | https://cgc.umn.edu/strain/JT23 |
| *ins-7(syb5424[ins-7::SL2::gfp::his-44]) IV.* | This paper | PHX5424 |
| *ins-33(syb5561[ins-33::SL2::gfp::his-44]) I.* | This paper | PHX5561 |
| *otEx8240[ges-1p::ins-7::tbb-2 3' UTR (8 ng/μL), ges-1p::gfp::his-44::tbb-2 3' UTR (3 ng/μL), unc-122p::mCherry (5 ng/μL), sheared OP50 genomic DNA (100 ng/μL)]* | This paper | OH19249 |
| *otEx8241[ges-1p::ins-7::tbb-2 3' UTR (8 ng/μL), ges-1p::gfp::his-44::tbb-2 3' UTR (3 ng/μL), unc-122p::mCherry (5 ng/μL), sheared OP50 genomic DNA (100 ng/μL)]* | This paper | OH19250 |
| *otEx8326[ges-1p::ins-33::tbb-2 3' UTR (8 ng/μL), ges-1p::gfp::his-44::tbb-2 3' UTR (3 ng/μL), unc-122p::mCherry (5 ng/μL), sheared OP50 genomic DNA (100 ng/μL)]* | This paper | OH19326 |
| *otEx8327[ges-1p::ins-33::tbb-2 3' UTR (8 ng/μL), ges-1p::gfp::his-44::tbb-2 3' UTR (3 ng/μL), unc-122p::mCherry (5 ng/μL), sheared OP50 genomic DNA (100 ng/μL)]* | This paper | OH19327 |
| *otEx8328[ges-1p::ins-33::tbb-2 3' UTR (8 ng/μL), ges-1p::gfp::his-44::tbb-2 3' UTR (3 ng/μL), unc-122p::mCherry (5 ng/μL), sheared OP50 genomic DNA (100 ng/μL)]* | This paper | OH19328 |
| *aex-1(ot1543[aex-1::SL2::gfp::his-44]) I.* | This paper | OH19333 |
| *aex-4(ot1530[aex-4::SL2::gfp::his-44]) X.* | This paper | OH19278 |
| *aex-5(ot1532[aex-5::SL2::gfp::his-44]) I.* | This paper | OH19280 |
| *otEx8059[ges-1p::ins-1::tagRFP-T::SL2::gfp::his-44::tbb-2 3' UTR (20 ng/μL), inx-6prom18::tagRFP-T::unc-54 3' UTR (8 ng/μL), sheared OP50 gDNA (100 ng/μL)]* | This paper | OH18447 |
| *daf-16(ot853[daf-16::mNG::AID]) I; otSi2[ges-1p::TIR1(F79G)::mRuby::unc-54 3' UTR *ieSi61] II; daf-2(e1370) III; otEx8220[ges-1p::ins-1::tagRFP-T::SL2::ebfp2::his-44::tbb-2 3' UTR (2 ng/μL), inx-6prom18::tagRFP-T::unc-54 3' UTR (8 ng/μL), sheared OP50 gDNA (100 ng/μL)]* | This paper | OH19091 |

| | | |
|---|---|---|
| *otIs904[ges-1p::ins-1::tagRFP-T::SL2::gfp::his-44::tbb-2 3' UTR (2 ng/µL), inx-6prom18::tagRFP-T::unc-54 3' UTR (8 ng/µL), GeneRuler 1 kb plus DNA ladder (100 ng/µL)] V.* | This paper | OH18707 |
| *aex-1(ot1357) I; otIs904[ges-1p::ins-1::tagRFP-T::SL2::gfp::his-44::tbb-2 3' UTR (2 ng/µL), inx-6prom18::tagRFP-T::unc-54 3' UTR (8 ng/µL), GeneRuler 1 kb plus DNA ladder (100 ng/µL)] V.* | This paper | OH18861 |
| *aex-1(ot1357) I.* | This paper | OH18501 |
| *nlp-40(tm4085) I.* | National Bioresource Project, Japan | NM4394 |
| *daf-16(ot853[daf-16::mNG::AID]) I; otIs908[pha-4prom2::TIR1(F79G)::mTur2::tbb-2 3' UTR (8 ng/µL), unc-122p::mCherry::unc-54 3' UTR (5 ng/µL), GeneRuler 1 kb plus DNA ladder (100 ng/µL)] V.* | This paper | OH19078 |
| *aex-1(ot1464) daf-16(ot853[daf-16::mNG::AID]) I; otIs908[pha-4prom2::TIR1(F79G)::mTur2::tbb-2 3' UTR (8 ng/µL), unc-122p::mCherry::unc-54 3' UTR (5 ng/µL), GeneRuler 1 kb plus DNA ladder (100 ng/µL)] V.* | This paper | OH19173 |
| *daf-16(ot853[daf-16::mNG::AID]) I.* | PMID: 32550509 | OH14125 |
| *aex-1(ot1464) daf-16(ot853[daf-16::mNG::AID]) I.* | This paper | OH19020 |
| *ins-35(syb6236[ins-35::SL2::gfp::his-44]) V.* | This paper | PHX6236 |
| *ins-7(ot1427) IV.* | This paper | OH18835 |
| *ins-35(ot1443) V.* | This paper | OH18895 |
| *ins-7(ot1427) IV; ins-35(ot1443) V.* | This paper | OH19012 |
| *otEx8242[ges-1p::ins-35::tbb-2 3' UTR (8 ng/µL), ges-1p::gfp::his-44::tbb-2 3' UTR (3 ng/µL), unc-122p::mCherry (5 ng/µL), sheared OP50 genomic DNA (100 ng/µL)]* | This paper | OH19251 |
| *otEx8243[ges-1p::ins-35::tbb-2 3' UTR (8 ng/µL), ges-1p::gfp::his-44::tbb-2 3' UTR (3 ng/µL), unc-122p::mCherry (5 ng/µL), sheared OP50 genomic DNA (100 ng/µL)]* | This paper | OH19252 |
| *otEx8222[ges-1p::ins-7::tbb-2 3' UTR (8 ng/µL), ges-1p::ins-35::tbb-2 3' UTR (8 ng/µL), ges-1p::gfp::his-44::tbb-2 3' UTR (3 ng/µL), unc-122p::mCherry (5 ng/µL), sheared OP50 genomic DNA (100 ng/µL)]* | This paper | OH19105 |
| *otEx8223[ges-1p::ins-7::tbb-2 3' UTR (8 ng/µL), ges-1p::ins-35::tbb-2 3' UTR (8 ng/µL), ges-1p::gfp::his-44::tbb-2 3' UTR (3 ng/µL), unc-122p::mCherry (5 ng/µL), sheared OP50 genomic DNA (100 ng/µL)]* | This paper | OH19106 |
| *otEx8224[ges-1p::ins-7::tbb-2 3' UTR (8 ng/µL), ges-1p::ins-35::tbb-2 3' UTR (8 ng/µL), ges-1p::gfp::his-44::tbb-2 3' UTR (3 ng/µL), unc-122p::mCherry (5 ng/µL), sheared OP50 genomic DNA (100 ng/µL)]* | This paper | OH19107 |
| *daf-7(e1372) III; otEx8223[ges-1p::ins-7::tbb-2 3' UTR (8 ng/µL), ges-1p::ins-35::tbb-2 3' UTR (8 ng/µL), ges-1p::gfp::his-44::tbb-2 3' UTR (3 ng/µL), unc-122p::mCherry (5 ng/µL), sheared OP50 genomic DNA (100 ng/µL)]* | This paper | OH19747 |
| *otIs937[ceh-19prom2::daf-2(DN)::eBFP2::SL2::tagRFP-T::tbb-2 3' UTR (10 ng/µL), unc-122p::gfp::unc-54 3' UTR (5 ng/µL), GeneRuler 1 kb plus DNA ladder (100 ng/µL)] V.* | This paper | OH19735 |

| | | |
|---|---|---|
| *otIs942[tph-1prom5::daf-2(DN)::eBFP2::SL2::tagRFP-T::tbb-2 3' UTR (12 ng/μL), unc-122p::gfp::unc-54 3' UTR (5 ng/μL), GeneRuler 1 kb plus DNA ladder (100 ng/μL)] V.* | This paper | OH19767 |
| *ins-7(ot1427) IV; otIs942[tph-1prom5::daf-2(DN)::eBFP2::SL2::tagRFP-T::tbb-2 3' UTR (12 ng/μL), unc-122p::gfp::unc-54 3' UTR (5 ng/μL), GeneRuler 1 kb plus DNA ladder (100 ng/μL)] ins-35(ot1443) V.* | This paper | OH19809 |
| *tph-1(ot1545) II.* | This paper | OH19337 |
| *otIs927[ges-1p::nlp-40::tagRFP-T::SL2::gfp::his-44::tbb-2 3' UTR (2 ng/μL), inx-6prom18::tagRFP-T::unc-54 3' UTR (8 ng/μL), GeneRuler 1 kb plus DNA ladder (100 ng/μL)] V.* | This paper | OH19575 |
| Oligonucleotides | | |
| Alt-R CRISPR-Cas9 tracrRNA | Integrated DNA Technologies | Catalog No. 1072533 |
| crRNA and ssODN sequences, see Table S3 | This paper | N/A |
| Software and algorithms | | |
| Behavioral Observation Research Interactive Software (BORIS) version 8.23 | DOI: 10.1111/2041-210X.12584 | https://www.boris.unito.it |
| Galaxy web platform | PMID: 38769056 | https://usegalaxy.org |
| FastQC Galaxy version 0.73 | Babraham Bioinformatics Group | https://www.bioinformatics.babraham.ac.uk/projects/fastqc |
| Trimmomatic version 0.38 | PMID: 24695404 | https://github.com/usadellab/Trimmomatic |
| STAR version 2.7.8a | PMID: 23104886 | https://github.com/alexdobin/STAR/releases |
| featureCounts version 2.0.1 | PMID: 24227677 | https://bioconductor.org/packages/release/bioc/html/Rsubread.html |
| DESeq2 Galaxy version 2.11.40.7 | PMID: 25516281 | https://github.com/thelovelab/DESeq2 |
| heatmap2 version 3.0.1 | R ggplot2 package | https://ggplot2.tidyverse.org |
| Venny version 2.1.0 | Juan Carlos Oliveros | https://bioinfogp.cnb.csic.es/tools/venny |
| Venn Diagram Plotter version 1.5.5228 | Pacific Northwest National Laboratory | https://github.com/PNNL-Comp-Mass-Spec/Venn-Diagram-Plotter |
| ShinyGO version 0.76 | PMID: 31882993 | https://bioinformatics.sdstate.edu/go |
| Fiji version 2.16.0 | PMID: 22743772 | https://imagej.net/software/fiji |
| G*Power version 3.1.9.2 | PMID: 19897823 | https://www.psychologie.hhu.de/arbeitsgruppen/allgemeine-psychologie-und-arbeitspsychologie/gpower |

| | | |
|---|---|---|
| R version 4.4.2 | The R Project for Statistical Computing | https://www.r-project.org |
| GraphPad Prism version 10.4.1 | GraphPad | https://www.graphpad.com |
| Other | | |
| Kontes Pellet Pestle Motor | Fisher Scientific | Catalog No. K749540-0000 |
